# Ketone body β-hydroxybutyrate restores neuronal Tau proteostasis via ketolysis-independent mechanism

**DOI:** 10.64898/2026.01.30.702936

**Authors:** Leigh-Ana M. Rossitto, Jamie Foo, Julia M. Di Silvestri, Leanne Lehmann, Melina Brunelli, Yuhang Nie, Daniel Muñoz-Mayorga, Qiang Xiao, Anna Y. Dai-Liu, Suborno Jati, Meghan Rossi, Richard Daneman, Jeffery W. Kelly, John C. Newman, Samuel A. Myers, Xu Chen

## Abstract

Metabolic interventions that induce ketosis, including ketogenic diets, caloric restriction, intermittent fasting, and exercise, show promise in the treatment of Alzheimer’s disease (AD) and related tauopathies. β-hydroxybutyrate (βHB), the primary ketone body produced during ketosis, reproduces key features of these metabolic interventions, but the molecular mechanism underlying its neuroprotective properties is not fully understood. Here, we demonstrate that a βHB precursor diet is sufficient to ameliorate Tau pathophysiology in a tauopathy mouse model. Furthermore, across *in vitro*, *ex vivo*, and *in vivo* models, we find that βHB enhances neuronal Tau proteostasis and reduces Tau aggregation and secretion. Importantly, these effects are independent of βHB’s oxidation for ATP production, as its ketolysis-resistant enantiomer reproduces these benefits, indicating that ketolysis is dispensable for these effects. Overall, these data position βHB as a novel therapeutic avenue for AD and tauopathy and elucidate a novel mechanism of action of metabolic interventions in neurodegenerative disease.

## INTRODUCTION

Alzheimer’s disease (AD) and related tauopathies are marked by brain accumulation of pathological Tau aggregates and progressive neurodegeneration, coinciding with metabolic changes such as glucose hypometabolism, lipid droplet accumulation, and elevated oxidative stress.^1–5^ At the organismal level, AD is associated with systemic metabolic dysfunction, and metabolic syndromes such as Type 2 diabetes, obesity, and hypertension increase AD risk.^6–10^ At the cellular level, single-nucleus RNA sequencing (snRNA-seq) of human brains reveals early and cell type-specific dysregulation of energy metabolism genes, particularly in neurons, supporting the concept that early neuronal energy crisis is a driver of disease progression.^11,12^ Together, these findings contribute to a growing interest in metabolism-based therapies for AD.^4^

Metabolic interventions including ketogenic diets, caloric restriction, intermittent fasting, time-restricted feeding, and exercise have shown promise in ameliorating cognitive impairments and neuropathology in AD models and patients.^13–24^ However, their translational potential is marred by variable efficacy, poor compliance in elderly and cognitively-impaired populations, caregiver burden, and adverse side effects.^25–29^ A shared molecular feature of these interventions is the induction of ketosis, during which *R*-β-hydroxybutyrate (βHB) is the primary circulating ketone body.^30^ While AD brains become insensitive to glucose, they maintain sensitivity to ketone bodies including *R-*βHB and increase βHB uptake from the blood, positioning βHB as a dietary mimetic for metabolic interventions.^31^ Previous studies demonstrate that βHB supplementation in the absence of complete dietary intervention reduces amyloid and Tau pathology and improves cognition in mouse models of AD, although the underlying molecular mechanism remains elusive.^17,32–36^

During ketolysis, *R-*βHB is oxidized by the enantiomer-specific enzyme *D-*beta-hydroxybutyrate dehydrogenase (BDH1) to acetoacetate (AcAc) for the production of acetyl-CoA and ATP. This process has several downstream metabolic and signaling consequences, including provision of alternative energy, modulation of sirtuin activity via cytoplasmic NAD+ sparing, and augmentation of the favorability of protein acetylation.^37–40^ Beyond ketolysis, βHB has emerged as a metabolite with a diverse repertoire of direct signaling functions, including G protein-coupled receptor binding, inhibition of Class I and IIa histone deacetylase complexes, inhibition of potassium efflux, βHBylation, and formation of βHB-amino acids.^40–46^ βHB’s dual ketolysis-dependent and - independent capacity suggests that its neuroprotective nature may extend beyond its catabolism for energy.

Here, we show that βHB alone is sufficient to ameliorate AD-relevant Tau pathophysiology in diverse *in vivo*, *ex vivo*, and *in vitro* model systems of tauopathy. Then, using *in vitro* and biochemical assays in combination with systemic approaches including snRNA sequencing and mass spectrometry-based Tau interactomics, we mechanistically demonstrate that βHB modulates Tau neuronal proteostasis, independent of ketolysis. These results elucidate a novel mechanism of action of metabolic interventions in tauopathy and poise the ketolysis-independent properties of βHB as novel therapeutic avenues for AD.

## RESULTS

### Therapeutic treatment of PS19 mice with a ketone ester diet ameliorates pathological Tau accumulation

Human mutant Tau^1N4R-P301S^ transgenic mice (herein PS19, hTau+ mice) develop progressive Tau pathology beginning at 5-6 months-old (mo) until the terminal age of 9-11 mo.^47^ We fed male hTau+ and wild-type (WT) littermate mice with a ketone ester diet (C6×2-*R*-1,3-BD, a βHB precursor, hence KE) or a macro-controlled diet (CTL) starting at approximately 6 mo for 16 weeks until 9-10 mo (**Fig. 1A-C**). The KE diet provides sustained low levels of βHB in a salt-free form through slower metabolism of the ketone ester (**Fig. S1A-C**).^48^ In line with previous reports, hTau+ mice lost approximately 20% of their original body weight during this time period.^49,50^ KE feeding prevented disease-associated weight loss in hTau+ mice without causing excess weight gain in WT mice (**Fig. 1D, S1D-F**). Mice on the KE diet consumed about the same number of kilocalories per gram of body weight as mice on the CTL diet within each genotype (**Fig. S1G-I**).

**Figure 1:**
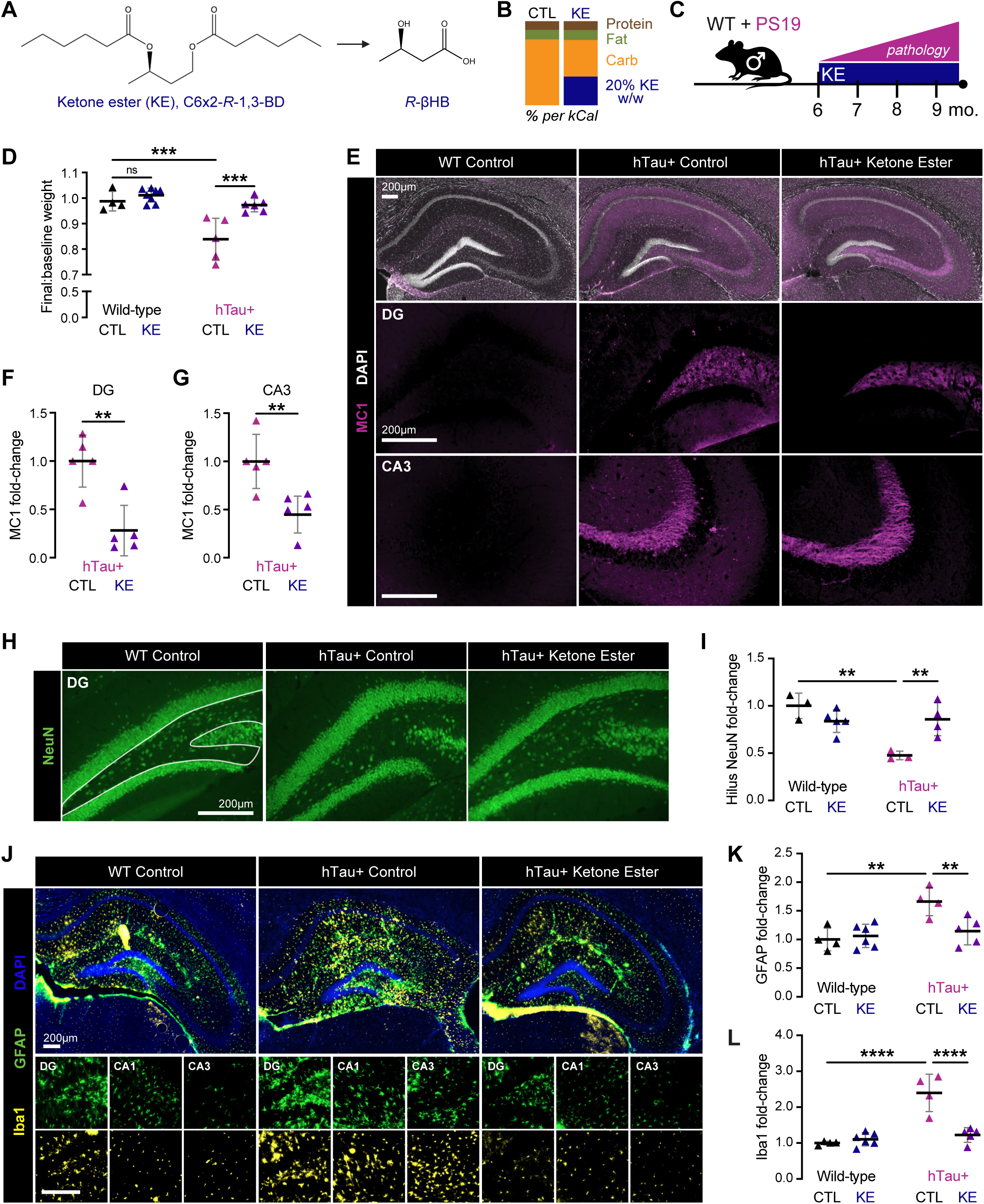
Ketone ester diet ameliorates neuropathology in PS19 mice. **(A)** Molecular structures of ketone ester C6×2-*R*-1,3-BD and *R-*βHB. **(B)** Macronutrient composition of the ketone ester (KE) diet (20% C6×2-*R*-1,3-BD w/w) and the macro-controlled (CTL) diet. **(C)** Schematic of ketone ester diet paradigm. **(D)** Ratio of end-point (week 16, approximately 9.5 mo) to baseline body weights of WT and hTau+ mice fed with CTL or KE diets. **(E-G)** Representative immunofluorescent images (E) and quantification (F-G) of dorsal hippocampi of WT and hTau+ fed with CTL or KE diet, stained for MC1 (magenta) and DAPI (white). Quantification represents the average of 3+ sections per mouse, normalized to the hTau+ CTL group. DG, dentate gyrus. Scale bars: 200 μm. **(H-I)** Representative immunofluorescent images (H) and quantification (I) of dorsal hippocampi of WT and hTau+ fed with CTL or KE diet, stained with anti-NeuN (mature neurons). Quantification represents NeuN+ area within the hilus of bregma-matched sections (white outline), normalized to the WT CTL group. Scale bar: 200 μm. **(J-L)** Representative immunofluorescent images (J) and quantifications (K-L) of dorsal hippocampi of WT and hTau+ fed with CTL or KE diet, stained with anti-GFAP (green, reactive astrocytes) and anti-Iba1 (yellow, microglia). Quantification represents the average value of 3+ sections per mouse, normalized to the WT CTL group. Scale bars: 200 μm. Data are represented as mean ± SD. Each point represents one mouse. ** p < 0.01, *** p < 0.001, **** p < 0.0001 by Welch’s t-test (F-G) and Šídák’s multiple comparisons test (D, I, K, and L).

To assess if the KE diet reduces disease-relevant, pathological Tau species, we performed immunofluorescent staining with the MC1 antibody, which recognizes a disease-associated conformational epitope of Tau (residues 7-9 and 312-342). The KE diet reduced MC1+ area by more than half in the dentate gyrus (DG) and CA3 subregions of the hippocampi of hTau+ mice (**Fig. 1E-G**). To assess for neuronal loss, we performed staining for mature neurons with anti-NeuN (neuronal nuclei protein RBFOX3) antibody (**Fig. 1H**). While we did not observe global neuronal loss in the hTau+ CTL diet mice, there were fewer NeuN+ cells within the DG hilus compared to WT, which has been reported in other mouse models of AD (**Fig. 1I**).^51^ KE feeding significantly rescued NeuN+ cell loss in DG hilus (**Fig. 1I**). Lastly, we assessed how the KE diet affected reactive astro- and microgliosis. As previously reported, hTau+ mice had a significant increase in GFAP+ and Iba1+ area in the hippocampus, in line with Tau pathology-associated neuroinflammation (**Fig. 1J-L**). Notably, the KE diet completely reverted GFAP+ and Iba1+ coverage to WT levels (**Fig. 1J-L**).

Next, to analyze levels of Tau species altered by a KE diet, we performed sequential biochemical fractionation of cortical tissue in a salt buffer (RAB), detergent buffer (RIPA), and 70% formic acid (FA) to assess highly soluble, less soluble, and highly insoluble Tau, respectively. We then probed for total Tau (HT7 for human Tau, hTau; DAKO for both human and mouse Tau, msTau) and phosphorylated Tau (pTau; AT8 for pTauS202/pT205; PHF1 for pTauS396/pS404) by immunoblotting. In the RAB and RIPA fractions, we did not observe significant reductions of total Tau in the KE-fed hTau+ mice (**Fig. S2A-G, Fig. 2A-F**). However, several pTau species were significantly decreased by the KE diet, specifically the 60 kDa pTau (by AT8) in the RIPA fraction, the 37 kDa (low molecular weight, LMW; cleaved) pTau (by PHF1) in the RAB and RIPA fractions, and the 120-180 kDa (high molecular weight, HMW; oligomeric) pTau (by AT8) in the RAB fraction (**Fig. S2A-G, Fig. 2A-F**). Most strikingly, in the insoluble (FA) fraction, we observed significant (∼50%) reductions by the KE diet in both total Tau (by HT7 and DAKO) and pTau (by PHF1 and AT8), as well as notable downshift of pTau band size (∼-5 kDa), in line with MC1 staining (**Fig. 2G-M**, **Fig 1E-G**).

**Figure 2:**
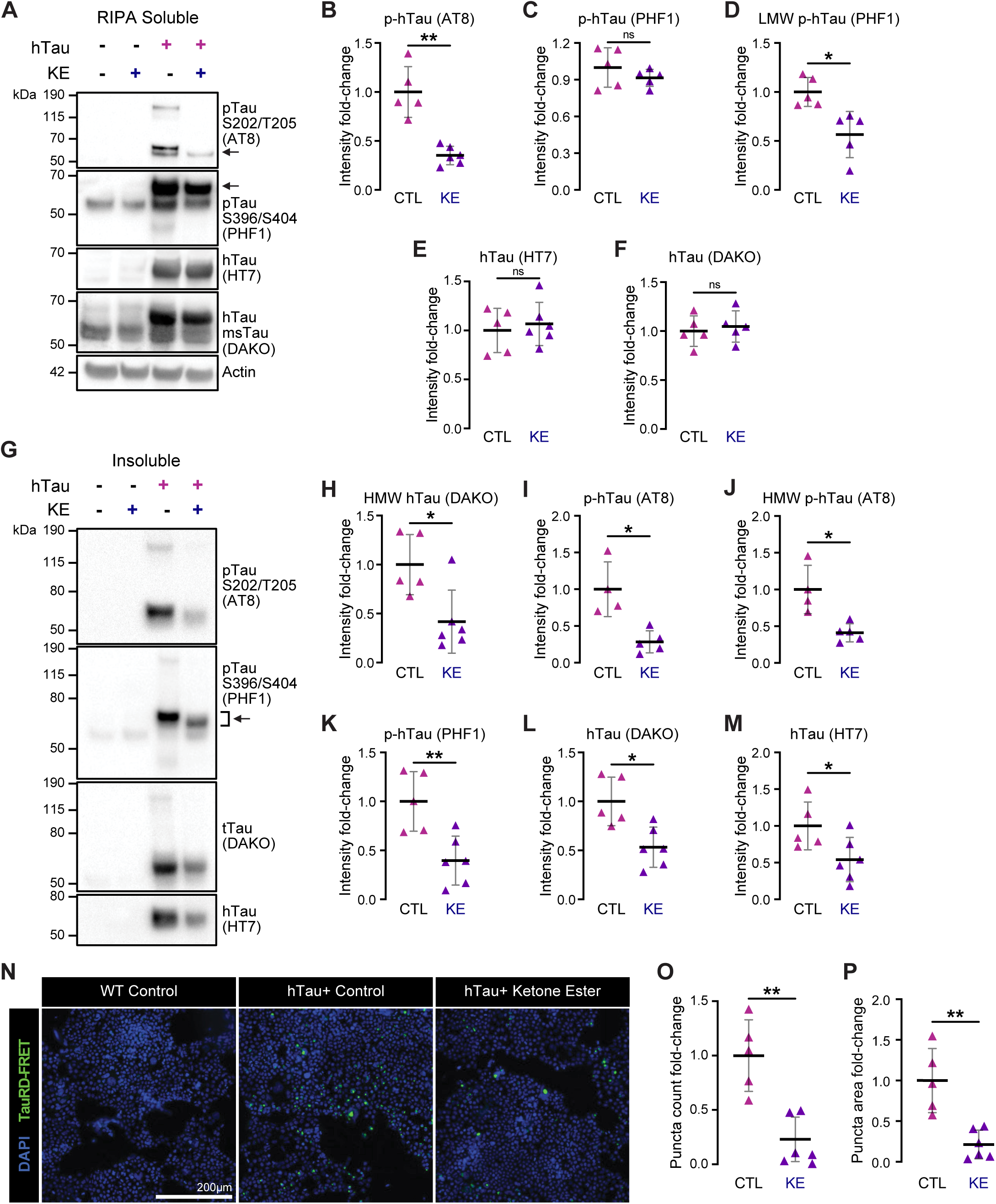
Ketone ester diet reduces levels of pathological Tau species, including hyperphosphorylated and insoluble Tau. **(A)** Representative Western blot showing the RIPA-soluble cortical fractions from WT and hTau+ mice fed with CTL or KE diets. Arrows indicate p-hTau band quantified for AT8 and PHF1. hTau, human Tau; msTau, mouse Tau. **(B-F)** Quantification of hTau from RIPA-soluble cortical fractions for pTau (B-D) and total hTau (E-F). LMW, low molecular weight (∼37 kDa). **(G)** Representative Western blot showing the formic acid-insoluble cortical fractions from WT and hTau+ mice fed with CTL or KE diets. Arrow and bracket indicate p-hTau band quantified for PHF1. Input was normalized by volume proportional to the initial amount (mg) of starting tissue. **(H-M)** Quantification of hTau from insoluble cortical fractions for pTau (H-K) and total hTau (L-M). HMW, high molecular weight (120-180 kDa). **(N-P)** Representative immunofluorescent images (N) and quantifications (O-P) of TauRD biosensor HEK293T cells seeded with PBS-soluble cortical fractions. Quantification represents the average value of two wells per mouse, four images per well. Scale bars: 200 μm. Data are represented as mean ± SD. Each point represents one mouse. Data were normalized to the hTau+ CTL group. * p < 0.05, ** p < 0.01, ns not significant by Welch’s t-test.

To assess the seeding capacity of Tau in brain lysates of hTau+ CTL and hTau+ KE mice, we performed a seeding assay with TauRD biosensor HEK293T cells.^52^ In the presence of pathological Tau seeds, CYP- and YFP-TauRD aggregate and generate FRET+ puncta. Commensurate with the results from our biochemical and immunofluorescent assessments, PBS-extracted cortical lysates from hTau+ KE mice induced fewer FRET+ puncta compared to hTau+ CTL mice (**Fig. 2N-P**). Immunoblotting showed a statistically insignificant trend of reduction in hTau and pTau levels in these PBS lysates **(Fig. S2H-K**). Taken together, these data demonstrate that KE diet feeding lowers pathological Tau species in hTau+ mouse brain, leading to less Tau phosphorylation, aggregation, and neurodegeneration.

### Hippocampal snRNA sequencing of ketone ester diet-fed PS19 mice highlights transcriptomic changes in excitatory neurons and glia

To begin to elucidate the molecular mechanisms underlying the benefit of the KE diet, we performed snRNA sequencing on flash-frozen hippocampi from hTau+ and WT mice fed with CTL and KE diets (n = 3/group). After filtering for high-quality singlets and merging the individual datasets, there were 183,015 total nuclei with comparable representation across groups (46,570, 45,836, 38,219, and 52,390 nuclei for WT CTL, WT KE, hTau+ CTL, and hTau+ KE groups, respectively) and a coverage of 28,363 genes for analysis. We identified 14 unique clusters of cells and used expression of canonical cell-type markers to classify nine clusters of neurons (seven excitatory, ExcN, and two inhibitory, InhN) and one cluster of each of the following: astrocytes (astros), microglia (micros), oligodendrocytes (oligos), oligodendrocyte precursor cells (OPCs), and vasculature-associated cells (vasc, **Fig. 3A, S3A**). We first compared cell-type distribution across the four groups (**Fig. 3B**). In line with previous snRNA-seq studies in AD mouse models and patient brains, hTau+ CTL diet mice had a significantly lower total proportion of ExcN compared to WT mice, especially ExcN1, resembling DG granule neurons (high in DG markers *Prox1* and *Calb1*) (**Fig. 3B, S3A**).^53,54^ hTau+ CTL mice also had significantly elevated proportions of microglia cells, consistent with increased Iba1+ immunohistochemistry (**Fig. 1L**), as well as oligodendrocytes (**Fig. 3B**). Remarkably, the KE diet significantly reverted the cell proportion alterations in the hTau+ mice (**Fig. 3B**). Additionally, hTau+ CTL mice showed a significant increase in *Gfap* and *Clec7a* expression in the astrocyte and microglia clusters, respectively, which was reduced by the KE diet at both the individual mouse and cell levels, consistent with immunofluorescent staining (**Fig. 3B, S3B-E**). Together, these data indicate a shift to reactive glia in the hTau+ CTL mice that is reverted to WT levels with the KE diet.

**Figure 3:**
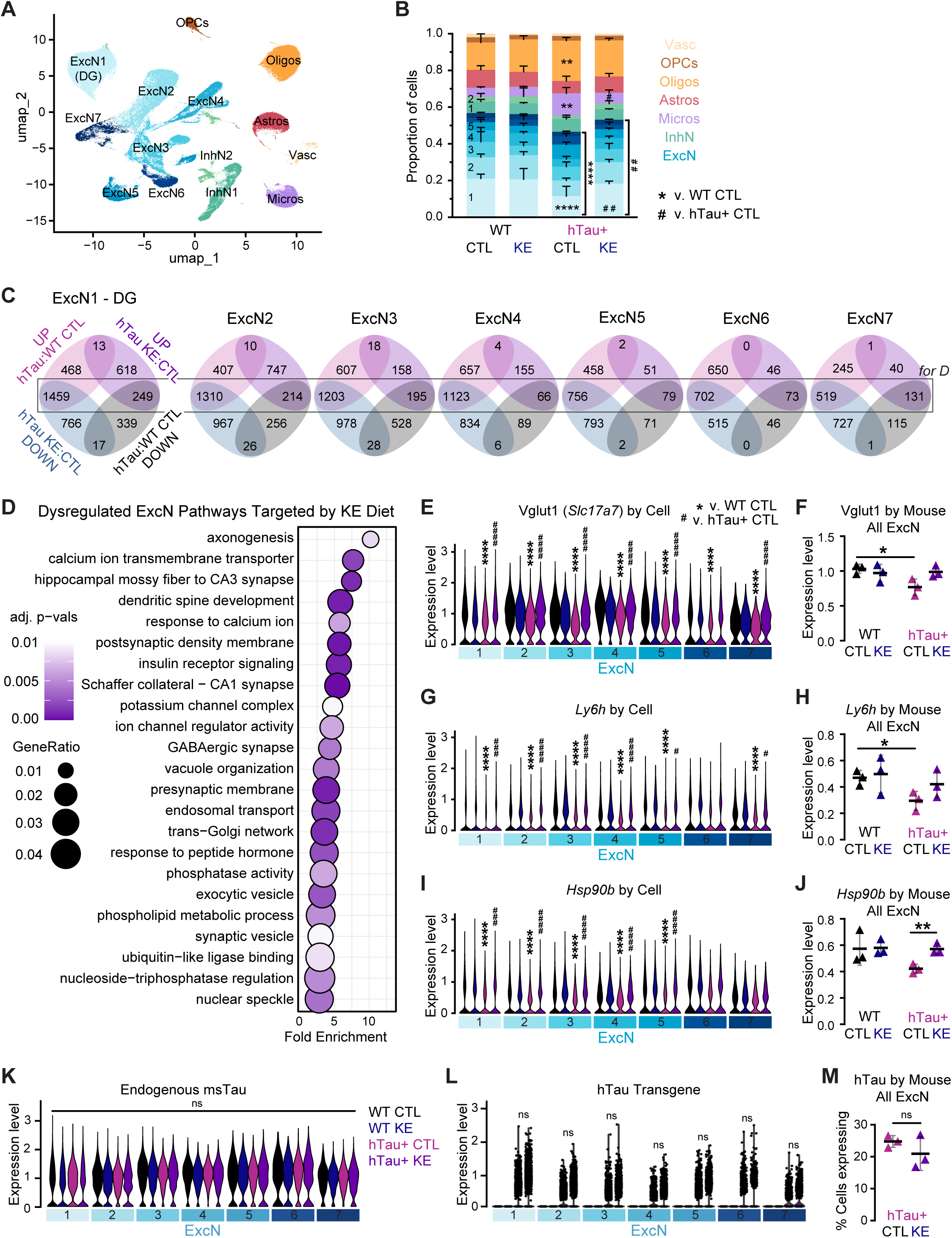
Hippocampal snRNA sequencing of ketone ester diet-fed PS19 mice highlights transcriptional changes in excitatory neurons and glia. **(A)** UMAP plot of 14 distinct cell clusters in the hippocampi of WT and hTau+ mice fed with CTL or KE diets (n = 3/group). ExcN, excitatory neurons; DG, dentate gyrus; InhN, inhibitory neurons; Micros, microglia; Astros, astrocytes; Oligos, oligodendrocytes; OPCs, oligodendrocyte precursor cells; and Vasc, vasculature-associated cells. **(B)** Bar graph depicting cell cluster proportion differences across groups. **(C)** Venn diagrams representing the number of significant DEGs in binary comparisons between hTau+ CTL v. WT CTL mice and hTau+ KE v. hTau+ CTL mice, by ExcN cluster. Top left (pink): genes upregulated in hTau+ CTL group compared to WT CTL group; top right (purple): genes upregulated in hTau+ KE group compared to hTau+ CTL group; bottom right (black): genes downregulated in hTau+ CTL group compared to WT CTL group; bottom left (blue): genes downregulated in hTau+ KE group compared to hTau+ CTL group. Boxed-in overlaps represent genes dysregulated in the hTau+ CTL mice that were reverted by the KE diet, used in 3D. **(D)** Gene ontology analysis of the list of dysregulated genes rescued in ≥4 ExcN clusters (see 3C). **(E-L)** Relative expression levels of Vglut1 (*Slc17a7*, E-F), *Ly6h* (G-H), *Hsp90b* (I-J), msTau (*Mapt*, K), and hTau (L) represented both at the single-cell level by ExcN cluster (E, G, I, K, L) and at the mouse level by pseudobulking (F, H, J). Data are presented by cluster in the following order: black, WT CTL; blue, WT KE; pink, hTau+ CTL; purple, hTau+ KE. **(M)** Percentage of total ExcNs expressing the hTau transgene, quantified at the mouse level. Data at the mouse level are represented as mean ± SD. * p < 0.05, ** p < 0.01, **** p < 0.0001 relative to WT CTL, and # p < 0.05, ### p < 0.001, #### p < 0.0001, ns not significant, relative to hTau+ CTL, unless comparisons are specified. DESeq2 test was used for expression level comparisons (E-L), Šídák’s multiple comparisons test for cell proportion (B), and percent expression comparisons (M).

Previous studies have implicated ExcNs at the center of the energy crisis in AD brains.^11,12,55^ We observed that *Oxct1*, the unidirectional ketolysis enzyme, is significantly increased in several ExcN clusters in hTau+ CTL compared to WT CTL mice, and MCT2 (*Slc16a7*), the βHB transporter with highest overall expression in our ExcN population, followed a similar pattern, suggesting a metabolic compensation in tauopathy neurons to ketone body utilization (**Fig. S3F-G**). To perform a deeper dive into network-level metabolic changes, we performed *in silico* metabolic flux predictions on micropooled ExcN transcriptomic profiles using Compass, a flux balance analysis method.^56^ Despite increased expression of ketone body utilization genes *Oxct1* and *Slc16a7*, Compass analysis revealed that hTau+ CTL neurons have decreased butanoate metabolic flux, suggesting impaired network-level ketone body metabolism (**Fig. S3H-J**). Beyond ketone bodies, hTau+ CTL ExcNs had discernable shifts in sugar and lipid metabolic flux, as revealed by principal component analysis in Compass space (**Fig. S3K-L**). Treatment with the KE diet restored hTau+ ExcN metabolic profiles to WT level.

Next, we performed differential expression analysis within each ExcN cluster, comparing differentially expressed genes (DEGs) between hTau+ CTL:WT CTL and hTau+ KE:hTau+ CTL, and overlaid the lists of DEGs between comparisons (**Fig. 3C**, **Table S1**). Strikingly, a large proportion of DEG transcripts found in hTau+ CTL mice were reverted in hTau+ KE mice (**Fig. 3C**). We next performed gene ontology (GO) analysis on the subset of DEGs that were reversed in ≥4 ExcN clusters, capturing 428 core genes increased in hTau+ CTL:WT CTL and decreased by the KE diet, and 44 core genes decreased in hTau+ CTL:WT CTL and restored by the KE diet (**Fig. 3C-D**). GO analysis revealed that the pathways targeted by the KE diet in ExcNs include metabolism, synaptic transmission and structure, the endomembrane system, and calcium signaling, among others, which are known dysregulated pathways in AD and tauopathy (**Fig. 3D**).

We observed transcriptomic evidence of dysregulated glutamatergic and cholinergic signaling. *Slc17a7*, which encodes for vesicular glutamate transporter Vglut1, a marker of glutamatergic synapses, was among the most significantly downregulated genes in hTau+ CTL mice at the cell and mouse levels and was rescued by the KE diet, suggesting restored excitatory synapse health in hTau+ mice (**Fig. 3E-F**). *Ly6h*, an antagonist of the hippocampal α7 nicotinic acetylcholine receptor (nAChR), was markedly reduced in hTau+ CTL mice and restored by the KE diet (**Fig. 3G-H**). Loss of Ly6h is linked to AD and neurodegeneration via dysregulated neuronal excitation and Tau hyperphosphorylation, and its antagonistic relationship with α7 nAChR was reflected across several ExcN clusters at the cell level (**Fig. S4A-B**).^57,58^ KE diet also reverted signatures of dysregulated neuronal proteostasis in the hTau+ CTL mice. Expression of *Hsp90* and other known Tau chaperones that escort aggregated Tau to both the proteasome and autolysosome was downregulated in hTau+ CTL mice and rescued by the KE diet (**Fig. 3I-J, S4C-H**). Lastly, hTau+ CTL mice showed increased neuronal C1q (*C1qa*–*c*) expression and corresponding elevation of microglial phagocytosis marker Gal-3 (*Lgals3*), a pattern consistent with microglia-mediated synaptic loss, which was also reverted by the KE diet (**Fig. S4I-J**).^59^

Overall, our snRNA-seq data shows that the KE diet rescues disease-associated pathways in hippocampal ExcNs, including metabolic and glutamatergic dysfunction. Notably, neither endogenous mouse *Mapt* (msTau) nor the human Tau (hTau) transgene was differentially expressed across clusters or treatment groups, indicating that the effects of the KE diet on Tau are likely by post-transcriptional mechanisms (**Fig. 3K-M**).

### βHB improves motor function and lifespan of Tau flies, driven by a ketolysis-independent mechanism

To dissect how βHB ameliorates tauopathy, we sought to disentangle actions that are dependent on ketolysis to generate energy (via acetyl-CoA and ATP production) from actions that are ketolysis-independent (via receptor binding, inhibition of Class I and IIa histone deacetylase complexes, βHBylation, etc.).^40^ We performed experiments comparing the canonical form of βHB, *R*-βHB, against its ketolysis-resistant enantiomer, *S*-βHB. *R*-βHB is the endogenous circulating metabolite produced by fatty acid oxidation and ketogenesis, oxidized to AcAc by the enantiomer-specific, rate-limiting enzyme BDH1, during ketolysis as the first step on the pathway to generate acetyl-CoA and ATP (**Fig. 4A**). *S*-βHB is not readily metabolized by BDH1, therefore is oxidized more slowly and incompletely, persisting longer in circulation.^60,61^ Accordingly, effects unique to βHB’s metabolism for energy should be relatively specific to *R*-βHB (v. *S*), whereas effects dependent on a ketolysis-independent property of βHB should be relatively specific to *S*-βHB or shared between *R*- and *S*-βHB.

**Figure 4:**
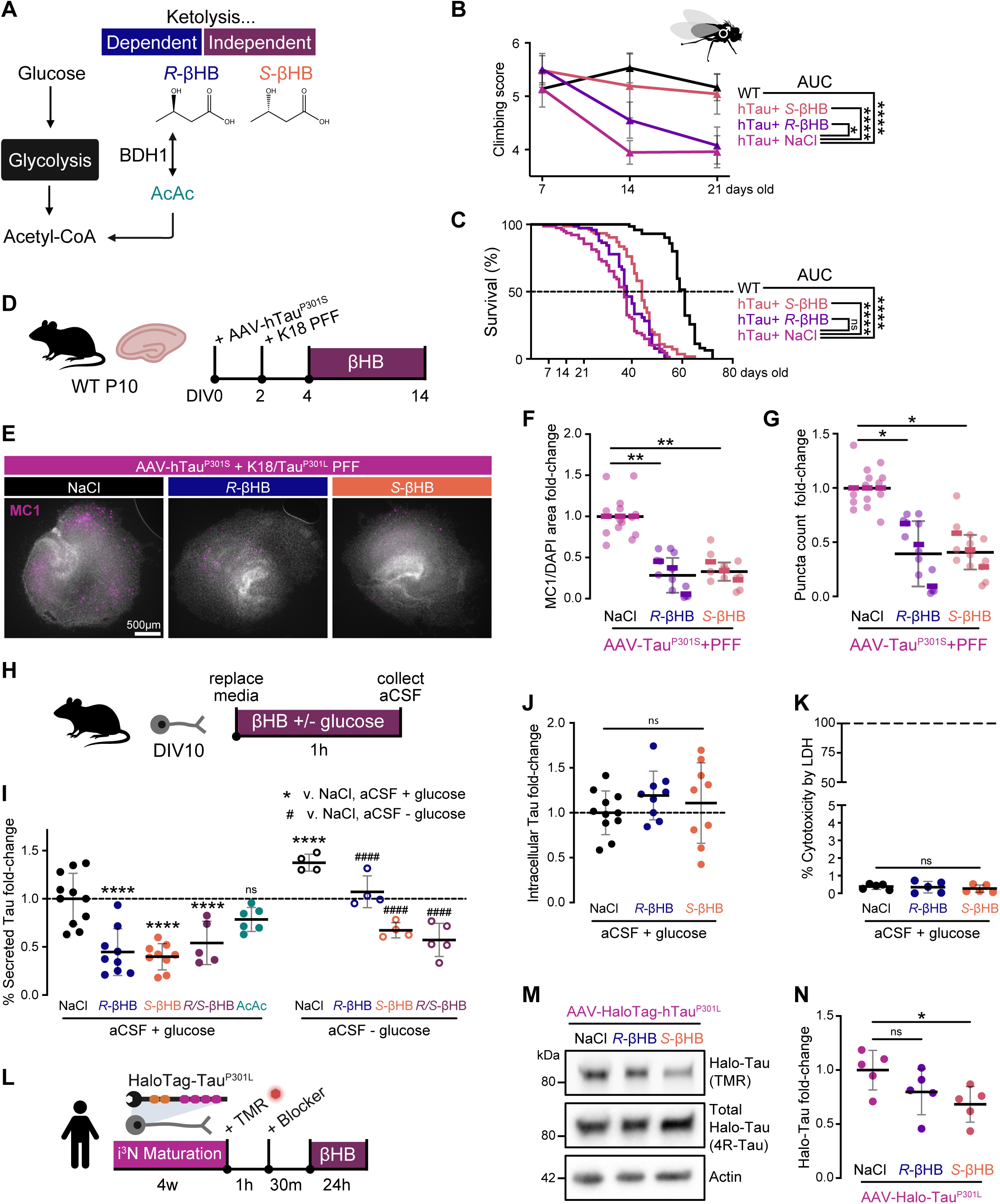
Ketolysis-independent properties of βHB promote Tau proteostasis and protect against Tau aggregation and neurotoxicity. **(A)** Schematic depicting enantiomer strategy for dissecting ketolysis-dependent and -independent features of βHB. AcAc, acetoacetate. **(B)** Climbing score for diet-fed WT and hTau+ flies at 7, 14, and 21 days old. Data represents the average scores of 4 vials per group from three trials. n = 15 flies/vial. AUC, area under the curve. **(C)** Survival curve for diet-fed WT and hTau+ flies. n = 20 flies/vial, 5 vials per group. **(D)** Schematic of organotypic brain slice culture (BSC) paradigm used in E-G. PFF, pre-formed fibrils. **(E-G)** Representative immunofluorescent images (E) and quantification (F, G) of BSC treated with AAV-Tau^P301S^, K18-Tau^P301L^ PFF, and βHB or NaCl for 10 days, stained for MC1 (magenta) and DAPI (white). Each point represents an individual brain slice, with slices in the same well stacked into one column. Thick, color-coded bars represent the well mean (n = 4-6 slices/well), and black bars represent the overall group mean ± SD (n = 3 wells, from separate batches). Each batch was normalized to the average of its respective control (NaCl) well. Scale bars: 500 μm. **(H)** Schematic of primary neuron Tau secretion paradigm for 4I. **(I)** Quantification of percent of Tau secreted into the conditioned media (CM) in primary neurons treated with various metabolites, with or without glucose, for 1 hr. Value calculated by the amount of Tau in the CM divided by the sum of the CM Tau and intracellular (lysate) Tau, and each point represents one independent well, normalized to the control (NaCl + glucose) well/s from its respective plate. Tau levels were measured using ELISA. aCSF, artificial cerebrospinal fluid. **(J)** Quantification of relative intracellular Tau levels in primary neurons, corresponding to 4I. Each point represents one independent well, normalized to the control (NaCl + glucose) well/s from its respective plate. **(K)** Lactate dehydrogenase (LDH) assay as a proxy for cytotoxicity. 100% represents the amount of LDH within the media upon complete cell lysis/death. **(L)** Schematic of HaloTag-Tau^P301L^ turnover assay in induced pluripotent stem cell-derived neurons (i^3^Ns). i^3^Ns do not express 4R-Tau normally, so an anti-4R-Tau antibody identifies total HaloTag-Tau. TMR, tetramethylrhodamine-conjugated Halo ligand. **(M-N)** Representative Western blot (M) and quantification (N) of Halo-Tau in-gel fluorescence. Each point represents one independent well, normalized to the average of the NaCl-treated values for its respective plate. Data are represented as mean ± SD. * p < 0.05, ** p < 0.01, **** p < 0.0001, ns not significant by AUC analysis (B, C), linear mixed-effects model with Tukey *post-hoc* test (F, G), and Dunnett’s multiple comparison test (J-K, N). For I, **** p < 0.0001, ns not significant compared to NaCl + glucose, #### p < 0.0001 compared to NaCl - glucose group by Šídák’s multiple comparisons test.

We first evaluated whether ketolysis is required for βHB’s protection against tauopathy in a *Drosophila melanogaster* model of tauopathy. These “Tau flies” overexpress human Tau pan-neuronally (*C155-Gal4>UAS-hTau^1.13^*), causing locomotor deficit and shortened lifespan.^62^ Adult Tau flies were cultured on food supplemented with 2 mM Na*R*-βHB, Na*S*-βHB, or NaCl (control). At Day 14, we evaluated motor deficits by a climbing assay, in which Tau flies begin to show locomotor deficit. Both *R*- and *S*-βHB-fed Tau flies performed significantly better than control Tau flies, with *S*-βHB-fed Tau flies performing as well as their WT counterparts (**Fig. 4B**). At Day 21 however, *R*-βHB-fed Tau flies no longer showed any improvement over the control, whereas *S*-βHB flies continued to show a complete rescue of locomotor deficit (**Fig. 4B**). We saw a similar pattern when assessing the impact of βHB supplementation on Tau fly lifespan: while the initial drop-off point for *R*-βHB-fed Tau flies is slightly later than NaCl-fed flies, there is no significant improvement to median or maximum lifespans nor overall area-under-the-curve (AUC), which represents the summation of number of days x flies alive (**Fig. 4C**). In contrast, *S*-βHB extended median lifespan from 37 to 44 days (WT being 61 days), maximum lifespan from 47 to 56 days (WT being 67 days), and significantly increased overall AUC (**Fig. 4C**). These results support that βHB ameliorates Tau-mediated neurotoxicity *in vivo* primarily through mechanisms independent of ketolysis.

### Ketolysis-independent properties of βHB promote Tau turnover and reduce Tau secretion in neurons

Next, we compared the effects of *R*- versus *S*-βHB on an *ex vivo* model of tauopathy, in which organotypic hippocampal slice cultures were infected with AAV9-hTau^P301S^ and seeded with K18/Tau^P301L^ fibrils to induce Tau accumulation (**Fig. 4D**). 10-day treatment with both *R*- and *S*-βHB significantly reduced the number and area of MC1+ Tau aggregates (**Fig. 4E-G**).

We then evaluated how βHB impacts neuronal Tau secretion, a critical first step in Tau propagation, in primary murine neurons (**Fig. 4H**). In the presence of glucose, 1 hr treatment with *R*-, *S*-, or racemic *R*-/*S*-βHB, reduced secreted Tau by ∼50%, whereas AcAc, the metabolite directly downstream of βHB in ketolysis, did not (**Fig. 4I**). This result supports the hypothesis that βHB inhibits Tau secretion via ketolysis-independent activity. No changes were observed in intracellular Tau levels (**Fig. 4J**), and no cytotoxicity was observed (**Fig. 4K**). We next tested whether removing glucose in the media, thereby forcing the neurons to utilize *R*-βHB as an alternative energy source, would amplify or blunt the effect of βHB on Tau secretion. Under glucose-deprived conditions, βHB remained effective in reducing Tau secretion, and the effect of *S*- and *R*-/*S*-βHB was more pronounced than *R*-βHB (**Fig. 4I**). We hypothesize that the dampened effect of *R*-βHB is likely due to its rapid oxidation as a fuel source under glucose starvation, causing a decrease in ketolysis-independent activity.

To assess how βHB affects Tau turnover in neurons and if ketolysis is required, we took advantage of a newly developed HaloTag-Tau^P301L^ tool to pulse-trace a subset of Tau in human induced pluripotent stem cell-derived neurons (i^3^Ns).^63^ After inducing HaloTag-Tau^P301L^ construct expression with doxycycline (dox), HaloTag-Tau was labeled for 1 hr with fluorescent tetramethylrhodamine (TMR) Halo ligand, followed by 24-hr treatment with *R*-βHB, *S*-βHB, or NaCl (**Fig. 4L**). In-gel fluorescence showed that *S*-βHB significantly reduced TMR-labeled HaloTag-Tau (**Fig. 4M-N**). Taken together, these *in vitro* data support that βHB reduces Tau secretion and promotes Tau turnover via ketolysis-independent action.

### βHB alters the neuronal Tau interactome, strengthening Tau’s interaction with endomembrane and synaptic systems, independent of ketolysis

Changes in Tau protein-protein interactions (PPI) underlie pathological events in AD including Tau aggregation, mislocalization, secretion, and propagation.^64^ To further understand how βHB alters Tau pathophysiology in neurons, we compared the PPI network of Tau under control and βHB-treated conditions in an i^3^N line engineered with a dox-inducible APEX2-N-Tau^2N4R^ (herein APEX-Tau) cassette.^64^ To account for Tau-unspecific PPIs, we utilized an isogenic APEX2-N-Tubulin α-1B (herein APEX-Tub) i^3^N line.^64^ Briefly, APEX-Tau and APEX-Tub i^3^Ns were differentiated and dox-treated to induce APEX2 cassette expression. After 30 min treatment with NaCl, *R*-βHB, or *S*-βHB, Tau- and tubulin-interacting proteins were then labeled with biotin, enriched by streptavidin pulldown, and analyzed by multiplexed tandem mass spectrometry (MS) (**Fig. 5A**). APEX-Tau proximal proteome alterations by βHB were determined by linear regression model to remove variance shared with APEX-Tub, which were considered background proteomic changes (see **Methods**).

**Figure 5:**
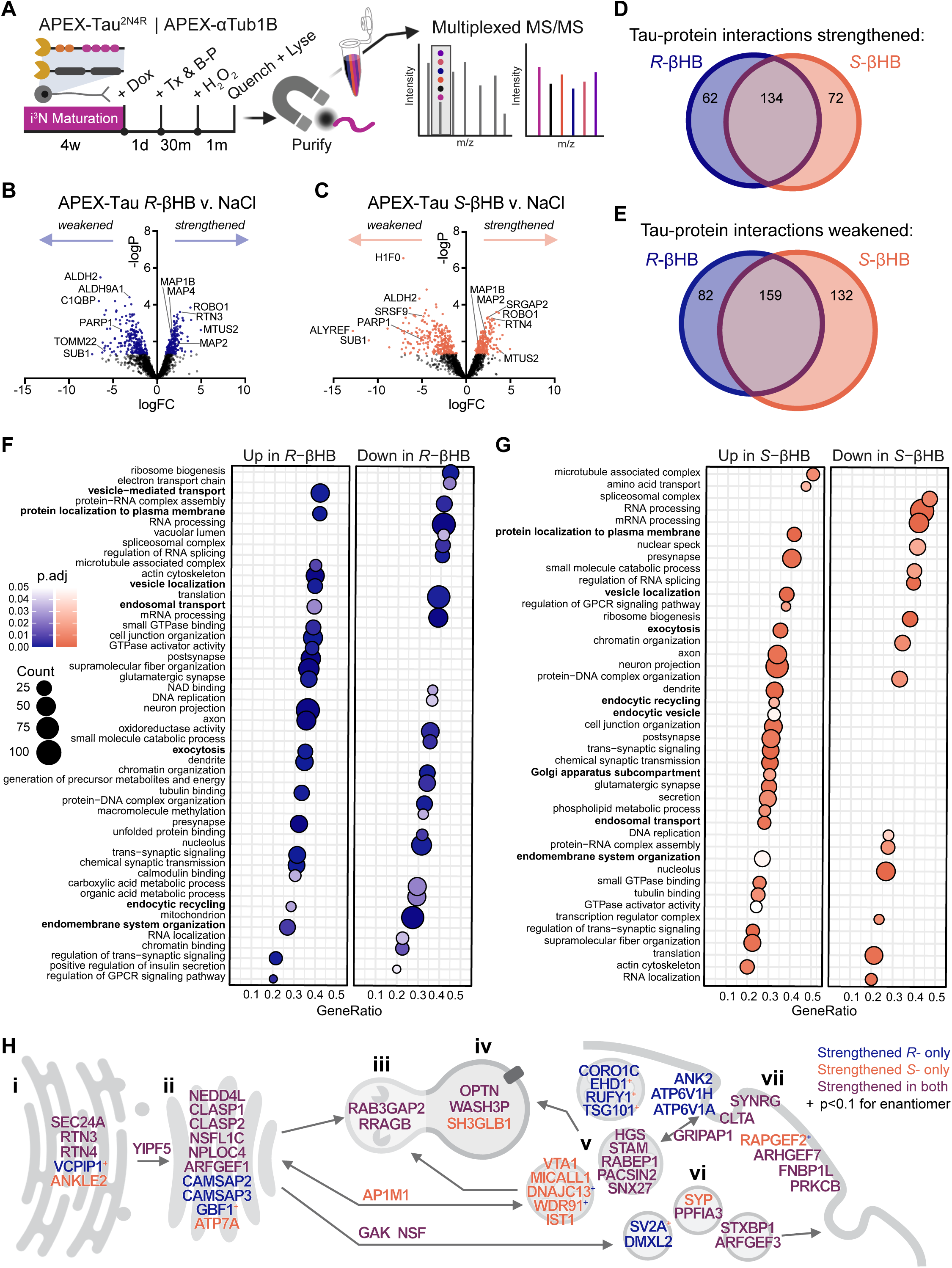
βHB alters neuronal Tau interactome, strengthening Tau’s interaction with endomembrane and synaptic systems. **(A)** Schematic overview of the APEX-Tau and APEX-Tub iPSC-derived neuron (i^3^N) constructs, labeling and treatment (tx) paradigm, and multiplexed mass spectrometry (MS) approach. Dox, doxycycline; B-P, biotin-phenol. **(B-C)** Volcano plots of binary comparisons between APEX-Tau NaCl v. *R*-βHB (B) or v. *S*-βHB (C), after linear regression model application to remove artifacts due to changes in the background proteome (see **Methods**). Each dot represents a single Tau interactor, and colored dots were significant by p < 0.05. **(D-E)** Venn diagrams depicting distinct and overlapping strengthened (D) and weakened (E) Tau interacting proteins by NaCl v. *R*-βHB or v. *S*-βHB. **(F-G)** Gene set enrichment analysis for *R*-βHB- (F) and *S*-βHB-associated Tau interactome. Input protein lists were ranked by score: -logP*FC. **(H)** Cartoon showing subcellular location of manually curated endomembrane system proteins whose interactions with Tau are strengthened with βHB. The color of the protein signifies the enantiomer of βHB that is significant. The “+” indicates that the change in interaction was significant in one enantiomer of βHB while the other enantiomer had a p-value < 0.1. i. Endoplasmic reticulum, ii. Golgi body, iii. Endolysosome, iv. Autophagosome, v. Endosomes, vi. Secretory vesicles, vii. Plasma membrane. P-values and FCs were determined by linear regression model to remove variance shared with APEX-Tub, representing artifactual interaction changes due to changes in the background proteome (see **Methods**). Significance cut-offs were defined as p < 0.05 and |FC| > 0.5 for D-H. FC, fold-change.

Both *R*- and *S*-βHB significantly altered the neuronal Tau interactome to a similar extent, as defined by number of differentially abundant interactors (i.e., “hits”): *R*-βHB and *S*-βHB significantly strengthening 196 and 206 Tau-specific protein interactions, while weakening 241 and 291 interactions, respectively (**Fig. 5B-C**). Among the hits, *R*- and *S*-βHB shared 134 strengthened and 159 weakened Tau-protein interactions (**Fig. 5D-E**). Gene set enrichment analysis (GSEA) revealed that both *R*- and *S*-βHB strengthened Tau-protein interactions with the endomembrane system (e.g., vesicle localization, endocytosis, and exocytosis) and synaptic transmission system (e.g., pre- and post-synapses and neuronal projections), and weakened PPIs related to nucleic acid metabolism, including the nucleus, ribosomes, and spliceosomes (**Fig. 5F-G**, **S5A**). Additionally, *R*-βHB, but not *S*-βHB, significantly decreased Tau’s interaction with mitochondrial and energy metabolism-related proteins (e.g., electron transport chain and NAD+ binding), suggesting that *R*-βHB uniquely modulates bioenergetics in Tau-bearing neurons (**Fig. 5F**).

Many of the differential interactors have reported roles in AD and tauopathy. βHB increased Tau’s interaction with SNAP91, a paralog of clathrin-mediated endocytosis protein PICALM, whose loss-of-function mutations are well-recognized genetic risk factors for sporadic AD (**Fig. S5B**).^65^ βHB also increased Tau’s interaction with components of ESCRT-0 (STAM and HGS) and ESCRT-I (TSG101), which are crucial for cargo sorting and trafficking to the autophagic-lysosomal system (**Fig. S5C-E**). Disruption of ESCRT machinery reportedly impairs Tau trafficking and increases Tau aggregation.^66,67^ ALYREF, an mRNA molecular chaperone and nuclear export adaptor whose knockout in *C. elegans* protected against Tau pathology and behavioral deficits, was the most significantly decreased Tau interactor by *S*-βHB by fold-change (p = 0.083 for *R*-βHB) (**Fig. S5F**).^68^ In conjunction with our previous results that show reverted synaptic transcriptome profile (**Fig. 3**) and improved Tau proteostasis (**Fig. 4**), these data support a model that βHB bolsters Tau’s roles in healthy neurons such as synaptic support and scaffolding (**Fig. S5A**), improves Tau proteostasis via entry into the endolysosomal system (**Fig. 5H**), and reduces Tau localization in organelles where aberrant, gain-of-function roles for Tau in disease have been reported (**Fig. 5F-G, S5F**).

### βHB improves Tau proteostasis via an OPTN-LC3B-dependent autolysosomal pathway

Optineurin (OPTN), a selective autophagy receptor in ubiquitin(Ub)-dependent mitophagy and aggrephagy, was identified as an APEX-Tau PPI whose interaction was significantly strengthened by both *R*- and *S*-βHB treatment (**Fig. 6A**).^69^ Previous studies show that loss of OPTN increases aggregated, hyperphosphorylated Tau and seed-induced Tau aggregation in human 4R-Tau i^3^Ns, whereas overexpression of OPTN *in vitro* and *in vivo* lowers Tau accumulation and aggregation.^70–74^ To validate that βHB strengthens OPTN-Tau interaction, we performed co-immunoprecipitation (co-IP) in HEK293T cells overexpressing OPTN-eGFP and mScarlet-Tau^WT^ or mScarlet-Tau^V337M^. 1 hr βHB (both *R*- and *S*-) treatment increased OPTN-eGFP co-IP with Tau via an anti-hTau antibody (HT7) (**Fig. 6B**). Next, as OPTN is Ub-dependent, we assessed Tau ubiquitination in HEK293T cells overexpressing mScarlet-Tau^V337M^ with 1-hr βHB treatment (**Fig. 6C**). Both *R*- and *S*-βHB increased Ub in the immunoprecipitated Tau fraction, i.e., Ub-Tau, consistent with increased *Ubb* expression observed in ExcN clusters by snRNA-seq (**Fig. 6C**, **S6A**). On the other hand, interaction of APEX-Tau with Ub-independent selective autophagy receptors NIPSNAP1, NIPSNAP2, PHB2, and FKBP8 were not changed significantly with βHB (**Table S2**).^75^ These data support a model that βHB increases Ub-Tau and enhances OPTN-dependent trafficking of Tau to the autophagic-lysosomal pathway, thereby promoting Tau degradation and reducing Tau propagation (**Fig. 6D**).

**Figure 6:**
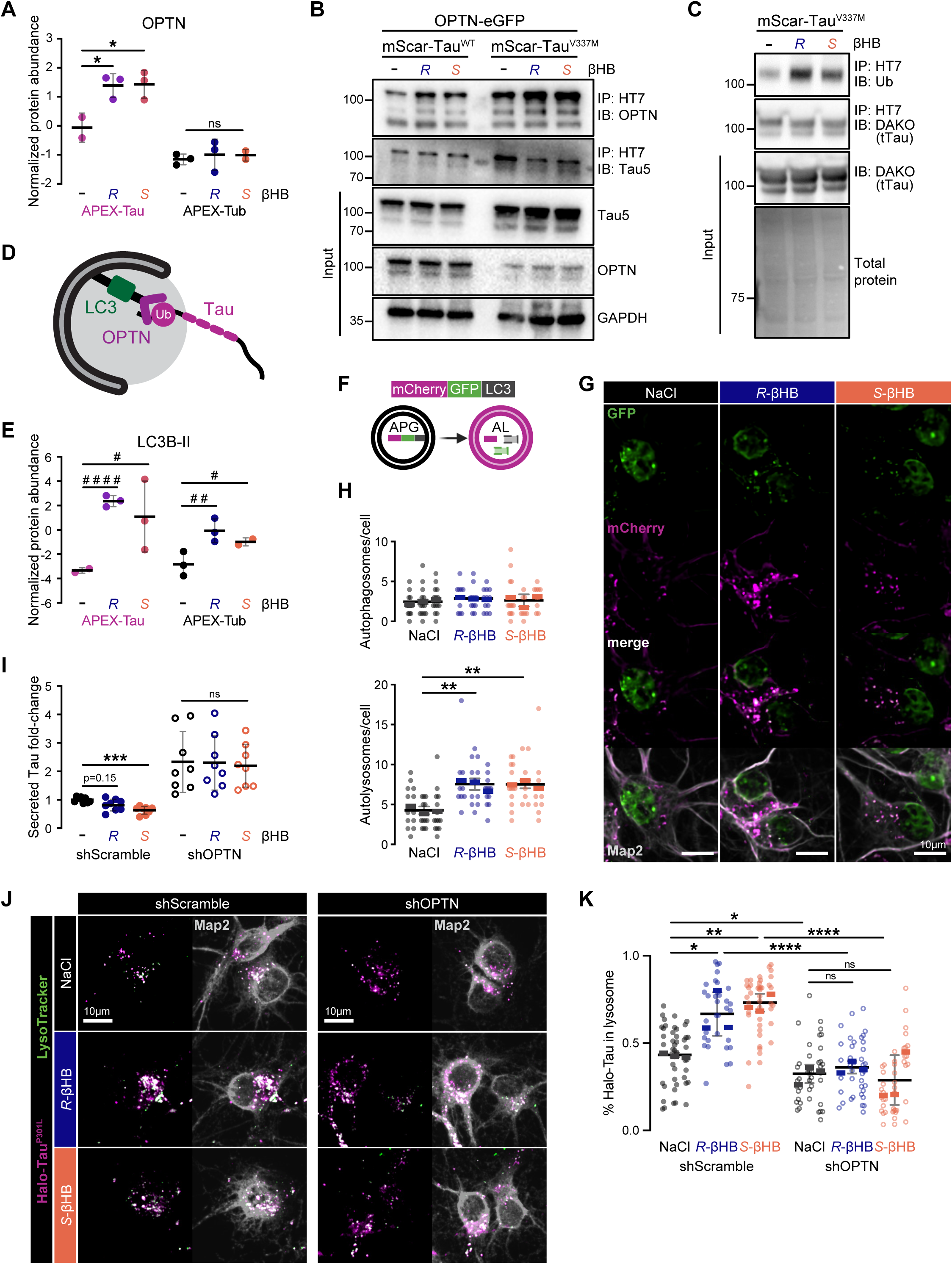
βHB improves Tau proteostasis via LC3B-OPTN axis. **(A)** Normalized protein abundances of Tau and Tubulin interactor OPTN from the interactome study. * p < 0.05 by linear regression model. **(B)** Representative Western blot of input lysate and co-IP with HT7 (hTau) in HEK293T cells overexpressing OPTN-eGFP and either mScarlet-Tau^WT^ or mScarlet-Tau^V337M^, treated with βHB. IP, immunoprecipitation; IB, immunoblot; Tau5, total Tau. **(C)** Representative Western blot of input lysate and IP with HT7 (hTau) in HEK293T cells overexpressing mScarlet-Tau^V337M^, treated with βHB for 1 hr. IP, immunoprecipitation; IB, immunoblot; DAKO, total Tau; Ub, ubiquitin. **(D)** Schematic of Ub-dependent recognition of Tau at the autophagosome membrane by the LC3B-adapter protein OPTN. **(E)** Normalized protein abundances of Tau and Tubulin interactor LC3B-II. # p < 0.05, ## p < 0.01, #### p < 0.0001 by pairwise limma test. **(F)** Graphic illustrating mCherry-GFP-LC3B construct. mCherry was pseudocolored to magenta. APG, autophagosome; AL, autolysosome. **(G-H)** Representative immunofluorescent images (G) and quantification of autophagic vesicles (H) in primary neurons transfected with lenti-mCherry-GFP-LC3B and treated with βHB for 1 hr. Magenta-only vesicles were counted as autolysosomes, and double-positive bright green and magenta vesicles (white) were counted as autophagosomes. Each point represents an individual Map2+ neuron, with cells in the same well stacked into one column. Thick, color-coded bars represent the well mean (n = 8-15 cells/well), and black bars represent the overall group mean ± SD (n = 3 wells/group, from separate batches). Scale bar: 10 μm. **(I)** Quantification of percent of Tau secreted into the conditioned media (CM) in primary neurons infected with either lenti-shScramble or lenti-shOPTN for >5 days and treated with βHB for 1 hr. Value calculated by Tau levels in the CM divided by the sum of the CM Tau and intracellular (lysate) Tau, measured by ELISA. Each point represents one independent well, normalized to the control (shScramble NaCl) wells from its respective plate. **(J-K)** Representative immunofluorescent images (J) and quantification of Halo-Tau^P301L^ (pink) co-localized with lysosomes (LysoTracker, green) in Map2+ neurons (K) co-transfected with lenti-Halo-Tau^P301L^ and either lenti-shScramble or lenti-shOPTN for 5 days then labeled and co-treated with βHB and LysoTracker for 1 hr. Values represent the area of Halo-Tau co-localized to lysosomes over the total Halo-Tau area within a single neuron. Each point represents an individual Map2+ neuron, with cells in the same well stacked into one column. Thick, color-coded bars represent the well mean (n = 10-20 cells/well), and black bars represent the overall group mean ± SD (n = 3 wells/group, from separate batches). Scale bar: 10 μm. Data for A, E, and I are represented as mean ± SD. * p < 0.05, ** p < 0.01, *** p < 0.001, **** p < 0.0001, ns not significant by linear mixed-effects model with Tukey *post-hoc* test (H, K) or by Šídák’s multiple comparisons test (I).

OPTN recruits ubiquitinated substrates to autophagosomes via binding to LC3B-II, the lipidated, membrane-bound form of LC3B that serves as a canonical marker for autophagosomes. A *post-hoc* pairwise comparison revealed that *R*- and *S*-βHB significantly increase Tau-LC3B-II interaction by 5.7- and 4.3-folds, respectively (**Fig. 6E**). *R*- and *S*-βHB also increased Tub-LC3B-II interaction, albeit by smaller magnitudes (**Fig. 6E**). It is well-documented that the ketogenic diet induces autophagy, and a recent study in primary neurons report that *R*-βHB promotes autophagic flux independent of glucose deprivation.^76–78^ To evaluate how *R*- and *S*-βHB affect neuronal autophagic flux, we transfected primary neurons with a dual-labeled fluorescent reporter, mCherry-GFP-LC3.^79^ In this system, autophagosomes (APGs) with a neutral pH appear white due to mCherry (pseudocolored magenta) and GFP fluorescence, whereas autolysosomes (ALs) with acidic pH are magenta due to quenching of the GFP fluorescence (**Fig. 6F**).^79^ 1 hr treatment of either *R*- or *S*-βHB increased the total number of autophagic vacuoles, exclusively driven by ALs (**Fig. 6G-H, S6B**).

To determine if βHB’s effects on Tau proteostasis are dependent at least in part by the OPTN-LC3B axis, we knocked down OPTN via lentivirally-delivered shRNA in primary neurons. shOPTN expression resulted in a near-complete loss of the OPTN protein (**Fig. S6C-D**). We performed a parallel Tau secretion assay comparing shOPTN-transduced neurons treated with and without βHB against their respective scramble control, shScr (**Fig. 6I, S6E**). Loss of OPTN increased Tau secretion at the baseline by about two-fold and abolished the benefit of either *R*- or *S*-βHB on reducing Tau secretion, without significantly altering intracellular Tau (**Fig. 6I, S6E**). Lastly, to visualize how βHB changes Tau recruitment to the lysosomes and whether it depends on OPTN, we co-transduced neurons with shOPTN and HaloTag-Tau^P301L^ lentiviruses, performed Halo-Tau labeling with fluorescent TMR for 1 hr, followed by 1 hr βHB and LysoTracker co-treatment (**Fig. 6J**). In the shScr group, both *R*- and *S*-βHB increased Halo-Tau co-localization with lysosomes (from 43% to ∼65-70% of the total Halo-Tau contained within lysosomes), and such increase was negated by OPTN depletion (**Fig. 6K**). Together, these results support a model that the ketolysis-independent activity of βHB promotes OPTN-dependent Tau recruitment to the autolysosomal pathway, thereby reducing Tau secretion and enhancing Tau degradation (**Fig. 6D**).

## DISCUSSION

The present study demonstrates that βHB, or a precursor in the form of a ketone ester, is sufficient to ameliorate AD-relevant Tau pathophysiology in diverse *in vivo*, *ex vivo*, and *in vitro* model systems, via a mechanism that does not require ketolysis to generate cellular energy and is dependent on LC3B-OPTN-mediated macroautophagy. Previously, Kashiwaya, et al. reported that a ketone ester diet can reduce pTau (T231) and improve cognitive endpoints in the 3xTg-AD model.^33^ To our knowledge, this is the first report that βHB protects against tauopathy in amyloid-independent models. Classically, βHB is elevated in states of nutrient and energy deprivation, such as fasting, starvation, and caloric restriction, as the organism switches towards lipid metabolism to provide alternative energy. However, the neuroprotective effect we and others observe with βHB supplementation without adopting a classic KD suggest that βHB supplementation could work as a ketomimetic, triggering cellular pathways that are activated by energy deprivation without commensurate cellular stress.^40^ Together, these results provide strong evidence that βHB supplementation is a promising intervention, adding to the growing body of work underscoring the utility of metabolic interventions in AD.^17,32–36^ To this aim, we offer our snRNA-seq and Tau interactome datasets as resources for the field to further our understanding of the molecular mechanisms driving the benefit of metabolic interventions in AD and tauopathy.

Here, we provide evidence that the ketolysis of *R*-βHB is, surprisingly, not required for its benefits in tauopathy. In *in vitro* (neuronal culture), *ex vivo* (BSC), and *in vivo* (fly) models, the ketolysis-resistant *S*-βHB fully recapitulates or even outperforms *R*-βHB in protecting against Tau aggregation, secretion, and neurotoxicity, supporting the idea that the benefits of βHB on Tau proteostasis are independent of βHB’s oxidation for energy. Indeed, in contexts that demand more ketolysis to generate alternative energy (e.g., longer timelines or glucose deprivation), the data suggest that the breakdown of *R*-βHB may blunt βHB’s benefits by leaving less βHB accessible for direct, ketolysis-independent cellular action. One caveat of using the *S*- and *R*-βHB enantiomers is that *S*-βHB, although metabolized differently and more slowly than *R*-βHB, may eventually be converted to *R*-βHB, AcAc, lipids, and CO_2_.^80^ Nevertheless, the different metabolic dynamics and route allow *S*-βHB to potentiate the ketolysis-independent action of βHB significantly longer than *R*-βHB, making it a useful tool for disentangling the two different aspects of βHB’s effects.^60,61^ Future studies should use BDH1-knockout models with *R*-βHB and AcAc to directly probe effects of βHB upstream and downstream of its metabolism, respectively.

Nonetheless, ketolysis-dependent features of βHB may still contribute to its benefit in tauopathy, despite them being dispensable for promoting neuronal Tau proteostasis, as in the present study. Generation of ATP via ketolysis helps circumvent glucose utilization, which is known to be impaired in tauopathy brains.^2,31^ Additionally, as *R*-βHB is metabolized to AcAc in the mitochondria whereas glycolysis occurs in the cytosol, cytosolic NAD+ is spared during *R*-βHB-driven ATP production, which can preserve metabolic flexibility and reduce glycolytic stress.^39^ Indeed, our snRNA-seq results suggest that KE diet (which provides *R*-βHB) reverses the aberrant metabolic shift in hTau+ neurons. Lastly, it is worth noting that intermediate metabolites of *R*-βHB metabolism, namely succinate and acetyl-CoA, can alter the favorability of proteins post-translational modifications, although the impacts of succinylation and acetylation on protein function are not completely known.^40^

In the present study, we identified an OPTN-LC3B autophagic-lysosomal axis as a key pathway mediating βHB’s effect on boosting Tau proteostasis. The autophagic-lysosomal pathway is not only pivotal for Tau degradation, but also critically involved in Tau trafficking and secretion.^67,81^ It has been previously reported that *R*-βHB is sufficient to activate autophagy in neurons.^78^ Here for the first time, we show that βHB induces autophagy via ketolysis-independent activity, mediated by Ub-dependent LC3B-adapter OPTN. OPTN has been reported to target soluble Tau towards autophagic-lysosomal degradation, and a few studies have implicated OPTN dysfunction in tauopathy.^70–74^ Our data suggest that βHB enhances OPTN recruitment of Ub-Tau to the autophagic-lysosomal pathway, leading to enhanced Tau degradation and less Tau secretion. How βHB regulates the OPTN-LC3B pathway is unclear. One possible upstream regulator of OPTN and autophagy is AMPK, which was found to be activated by βHB in various cell types.^78,82,83^ AMPK activates TBK1, which phosphorylates OPTN, thereby enhancing OPTN-LC3B binding affinity and autophagic flux.^84–87^ This axis represents a potential fasting-mimetic signaling activity of βHB underlying its Tau-lowering effect.

There are likely other metabolism-independent pathways mediating βHB’s neuroprotective effects in tauopathy. βHB can directly bind to two known G-protein-coupled receptors, HCAR2 and FFAR3, neither of which appear to be highly expressed in neurons, therefore their roles in regulating neuronal Tau proteostasis are unclear.^41,42,88^ These receptors are highly expressed on glia cells, especially microglia. Recent studies have linked HCAR2 and FFAR3 to antiseizure and anti-inflammatory effects of βHB.^44,89–91^ Future work should focus on elucidating cell-type specific roles of these βHB receptor pathways in glia, as well as cell non-autonomous effects in neuron-glia interactions. βHB is thought to be protective in epilepsy by improving excitatory:inhibitory imbalance, which is a shared pathophysiological feature in AD neurons.^92^ βHB has been shown to inhibit activity of neuronal vesicular glutamate transporter Vglut2 and increase GABA production, although conflicting evidence exists.^93–95^ In our snRNA-seq dataset, Vglut2 (*Slc17a6*) was most highly expressed in ExcN clusters 6 and 7, but there were no significant differences in expression across any of the groups (**Table S1**). On the other hand, we found Vglut1 (*Slc17a7*) expression decreased in ExcN of hTau+ mice, which was rescued by the KE diet. Exactly how βHB modulates neurotransmission warrants further study. βHB can alter gene transcription via inhibition of class I histone deacetylase complexes (HDACs 1, 2, 3, and 8) and via βHBylation of histones directly.^43,45,95^ We did observe a mild decrease of HDAC1 and HDAC8 in our snRNA-seq, along with transcript-level changes of vesicular transport and ubiquitin pathways in neurons, which could be partly responsible for the observed Tau proteostasis enhancement. Lastly, it is possible that Tau itself could be βHBylated, a post-translational modification on lysine residues, the implication of which remains to be elucidated.^36^

Current clinical trials are underway evaluating the potential of βHB precursors and other ketone body supplements in AD, other neurodegenerative diseases, and aging. While this study demonstrates that *S*-βHB may have more potent and consistent benefits than its metabolically active counterpart *R*-βHB, there are unique challenges in translating this finding to a therapeutic for patients, as *S*-βHB is not generated in significant quantities by endogenous ketogenesis and so cannot be provided to patients in the form of fatty acid precursor supplements. Potential translational avenues of the present study include the development and evaluation of BDH1-independent *S*-βHB precursors (e.g., *S*-1,3-butanediol) and the targeting of downstream Tau-modulatory pathways such as OPTN-LC3B-autophagy. Several current FDA-approved drugs including rapalogs and metformin are currently in clinical trials for MCI and AD, with diverse mechanisms of action including mTOR inhibition and AMPK/ULK1 activation. βHB-inducing interventions, alone or in combination with current autophagic-enhancing agents, may offer better efficacy and more desirable side effect profiles than current metabolic therapies.

### Limitations of the study

In our *in vivo* long-term supplementation, plasma βHB concentration was detected much lower than that tested in our acute *in vitro* experiments, although the levels are with previously published guidelines on nutritional ketosis.^96^ The relatively mild ketosis is not unexpected, because mice may adapt to βHB supplementation over a long feeding period. A second limitation is that our *in vivo* studies only utilized male mouse and fly models, whereas *in vitro* cultures and *ex vivo* slices were mixed-sex. Although it is common for studies on PS19 mice to use only male mice as they develop earlier and more consistent Tau pathology than females, future studies should evaluate sex differences in both the efficacy and the molecular mechanism of ketomimetic interventions. Lastly, clinical translation of βHB supplementation in mice may be limited by the difference in the macronutrient composition of standard rodent chow and human diets, in addition to the previously described challenge of synthesizing *S*-βHB-specific precursors.

## SUPPLEMENTAL FIGURES

**Figure S1, related to Figure 1:**
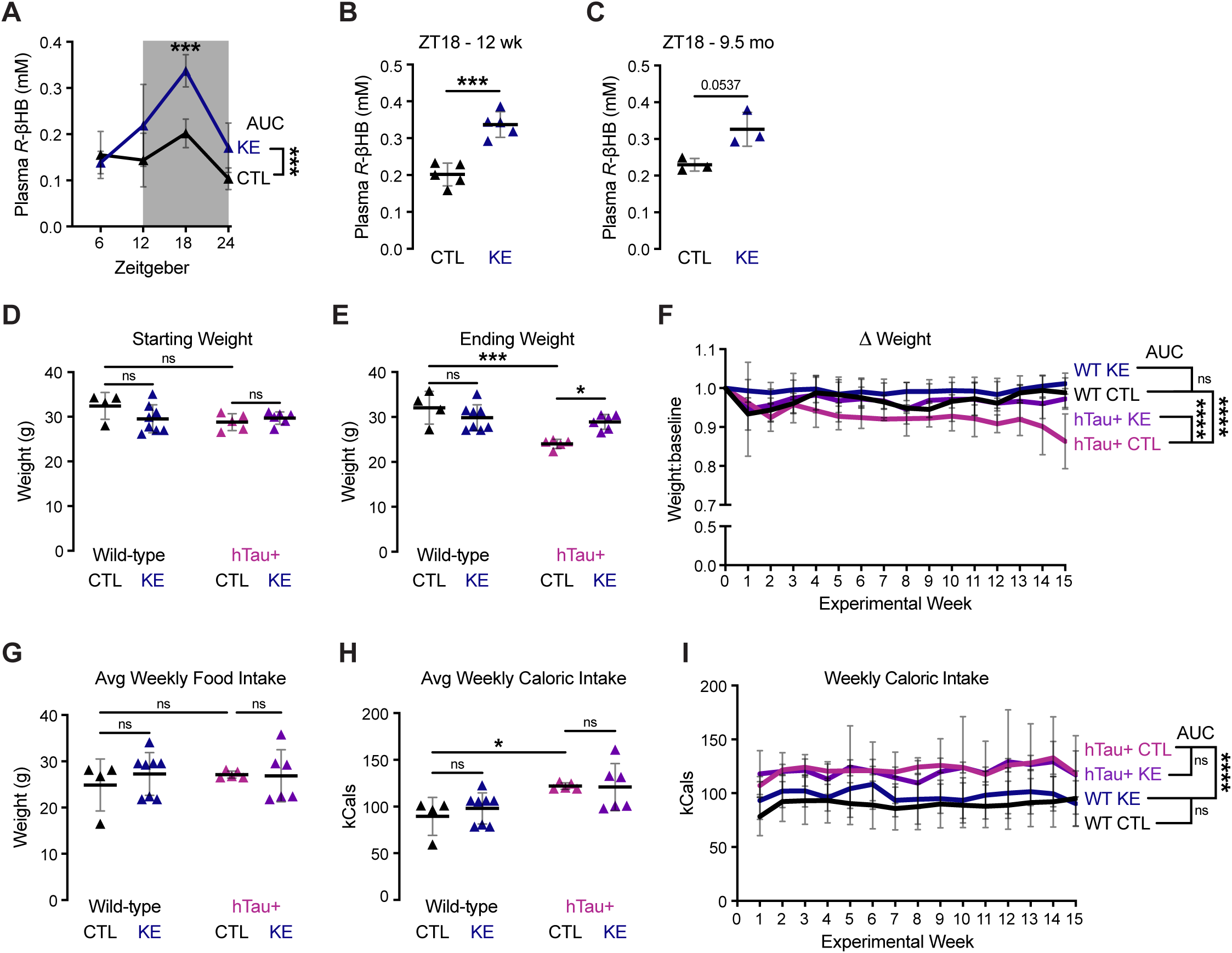
Ketone ester diet induces mild ketosis, and any weight or food intake differences from the diet are genotype-dependent. (A) Plasma *R-*βHB levels across Zeitgeber times (ZT) 6-24 for 12 week-old WT mice (C57B/6J) on the KE or CTL diets. AUC, area under the curve. (B) Plasma *R-*βHB levels at ZT18 at 12 weeks-old as in (A), showing individual mice. (C) Plasma *R-*βHB levels at ZT18 for 9.5 month-old (mo) CTL and KE diet fed mice. (D) Initial body weights of WT and hTau+ mice before administration of either diet, illustrating no baseline differences between the groups. (E) Ending body weights of WT and hTau+ mice at the end of the 16-week KE diet study. (F) Weight change compared to baseline by week for ketone ester diet study. **(G-I)** Average weekly food intake (G), average weekly kCals intake (H), and kCals intake by week (I). hTau+ mice consumed more chow regardless of diet. Data are represented as mean ± SD. * p < 0.05, ** p < 0.01, *** p < 0.001, **** p < 0.0001, ns not significant by Šídák’s multiple comparisons test (A, B, D, E, G, and H), Welch’s t-test (C), and AUC analysis (A, F, and I).

**Figure S2, related to Figure 2:**
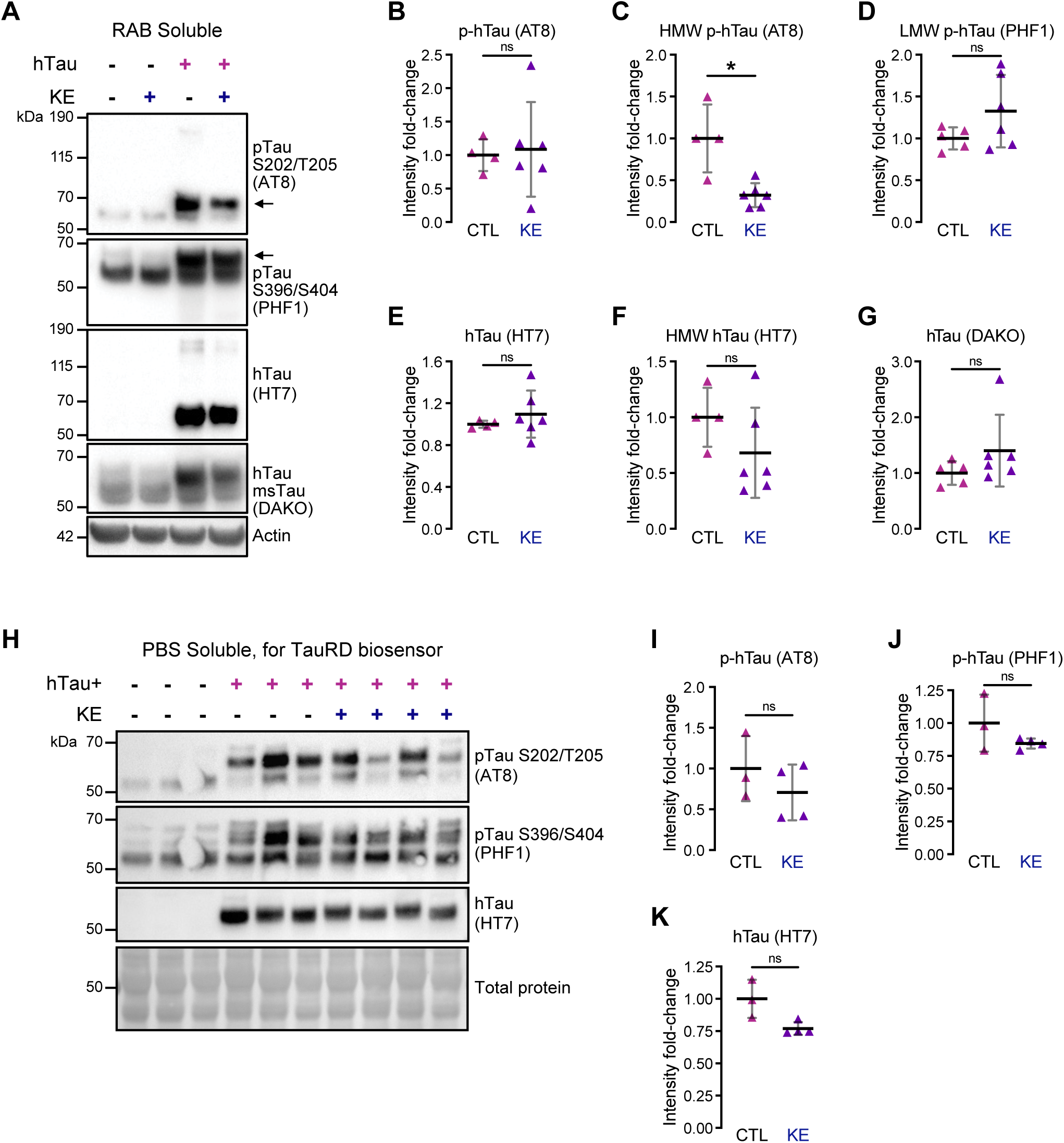
Ketone ester diet minimally changes highly soluble (RAB and PBS) species of Tau. **(A)** Representative Western blot showing RAB-soluble cortical fractions from WT and hTau+ mice fed with CTL or KE diets. Arrows indicate p-hTau band quantified for AT8 and PHF1. hTau, human Tau; msTau, mouse Tau. **(B-G)** Quantifications of p-hTau (B-D), hTau (E-F), and total Tau (G) in RAB-soluble cortical fractions. **(H)** Representative Western blot for Tau species in PBS-soluble cortical lysates from WT and hTau+ mice fed with CTL or KE diets. Lysates were used for a Tau seeding assay in HEK293T biosensor cells (Fig. 2N). **(I-K)** Quantifications of p-hTau (I-J) and total hTau (K) from PBS-soluble cortical lysates. Data are represented as mean ± SD. Each point represents one mouse. Data were normalized to the hTau+ CTL group. * p < 0.05, ns not significant by Welch’s t-test.

**Figure S3, related to Figure 3:**
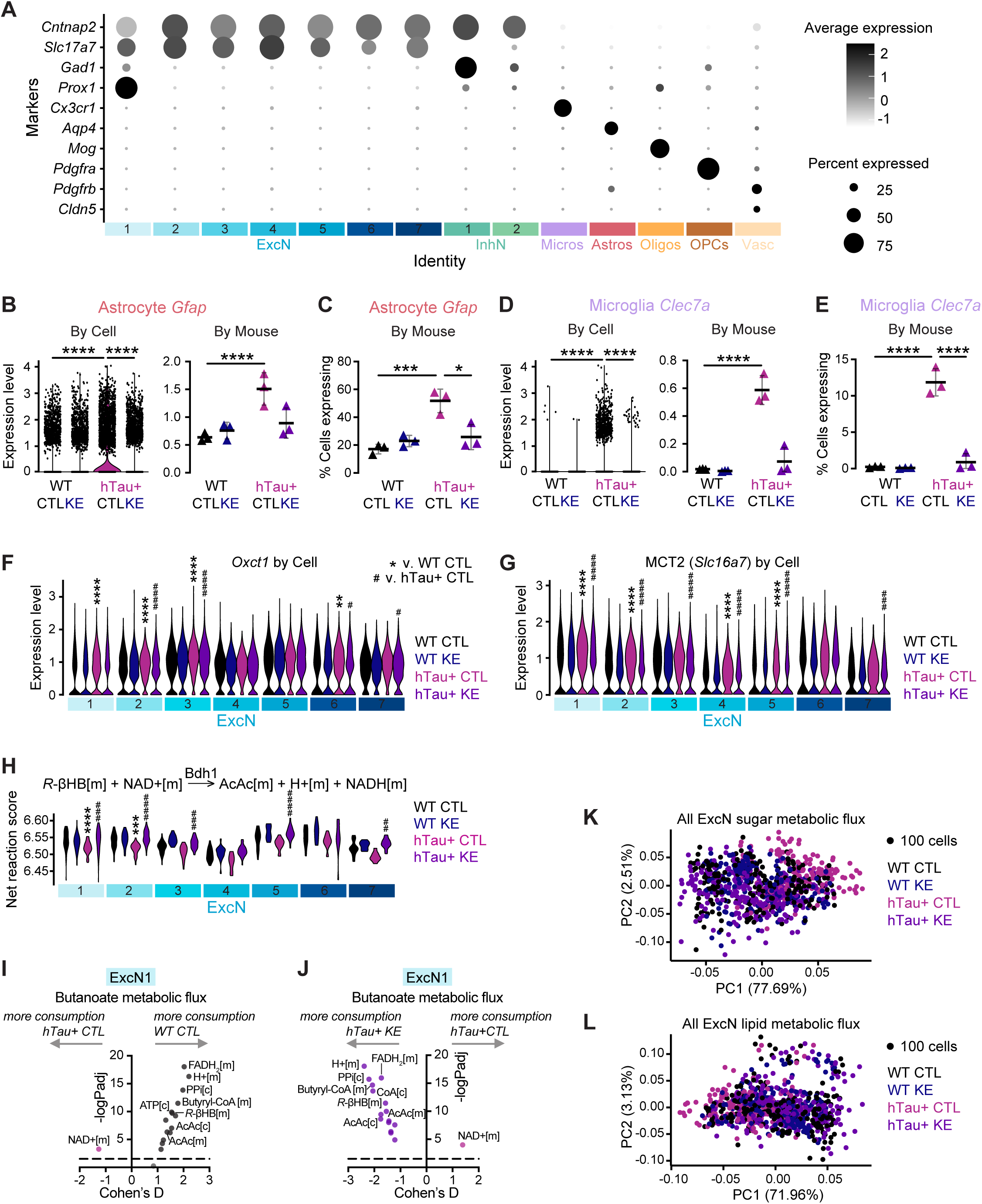
Hippocampal snRNA sequencing of KE diet-fed PS19 mice identifies 14 distinct cell types, with the KE diet reverting global cell-type distribution and metabolic profiles of hTau+ mice to WT levels. **(A)** Relative expression of canonical cell type markers in hippocampal snRNA seq experiment within diet study. 14 distinct clusters were identified. Circle saturation represents the normalized average expression of marker genes of all cells within a cluster, and size represents percent of cells within a cluster expressing a given marker gene. ExcN, excitatory neurons; InhN, inhibitory neurons; Micros, microglia; Astros, astrocytes; Oligos, oligodendrocytes; OPCs, oligodendrocyte precursor cells; and Vasc, vasculature-associated cells. **(B-C)** Cell- and mouse-level relative expression of *Gfap* (B), a reactive astrocytic marker, in the astrocyte cluster. The KE diet ameliorates elevated expression levels and proportion of astrocytes per mouse expressing *Gfap* (C) in hTau+ mice. **(D-E)** Cell- and mouse-level relative expression of *Clec7a* (D), a reactive microglial marker, in the microglia cluster. The KE diet ameliorates elevated expression levels and proportion of microglia per mouse expressing *Clec7a* (E) in hTau+ mice. **(F-G)** Relative expression levels of *Oxct1* (F) and *Slc16a7* (MCT2) (G), genes involved in ketone body utilization, quantified at the single-cell level by ExcN cluster. **(H)** Violin plot representing net (positive - negative) reaction scores for Bdh1-dependent ketolysis at the micropooled cell level by ExcN cluster. A larger net reaction score indicates more positive flux in the indicated reaction direction. [m] mitochondria. **(I-J)** Volcano plots of binary comparisons between hTau+ CTL v. WT CTL (I) or v. hTau+ KE (J) on micropooled ExcN1 net reaction scores for butanoate subsystem metabolites. Colored dots were significant by adjusted p < 0.05. [m] mitochondria, [c] cytosol. **(K-L)** Principal component (PC) analysis of Compass results for all ExcNs for sugar (K) and lipid (L) metabolism pathways. A single dot represents 100 micropooled cells within the same ExcN cluster and mouse. Data at the mouse level are represented as mean ± SD. ** p < 0.01, *** p < 0.001, **** p < 0.0001 relative to WT CTL, and # p < 0.05, ## p < 0.01, ### p < 0.001, #### p < 0.0001 relative to hTau+ CTL, unless comparisons are specified, by DESeq2 test (B, D, F-G), Šídák’s multiple comparisons test (C, E), and Welch’s t-test with Bonferroni’s correction (H-J).

**Figure S4, related to Figure 3:**
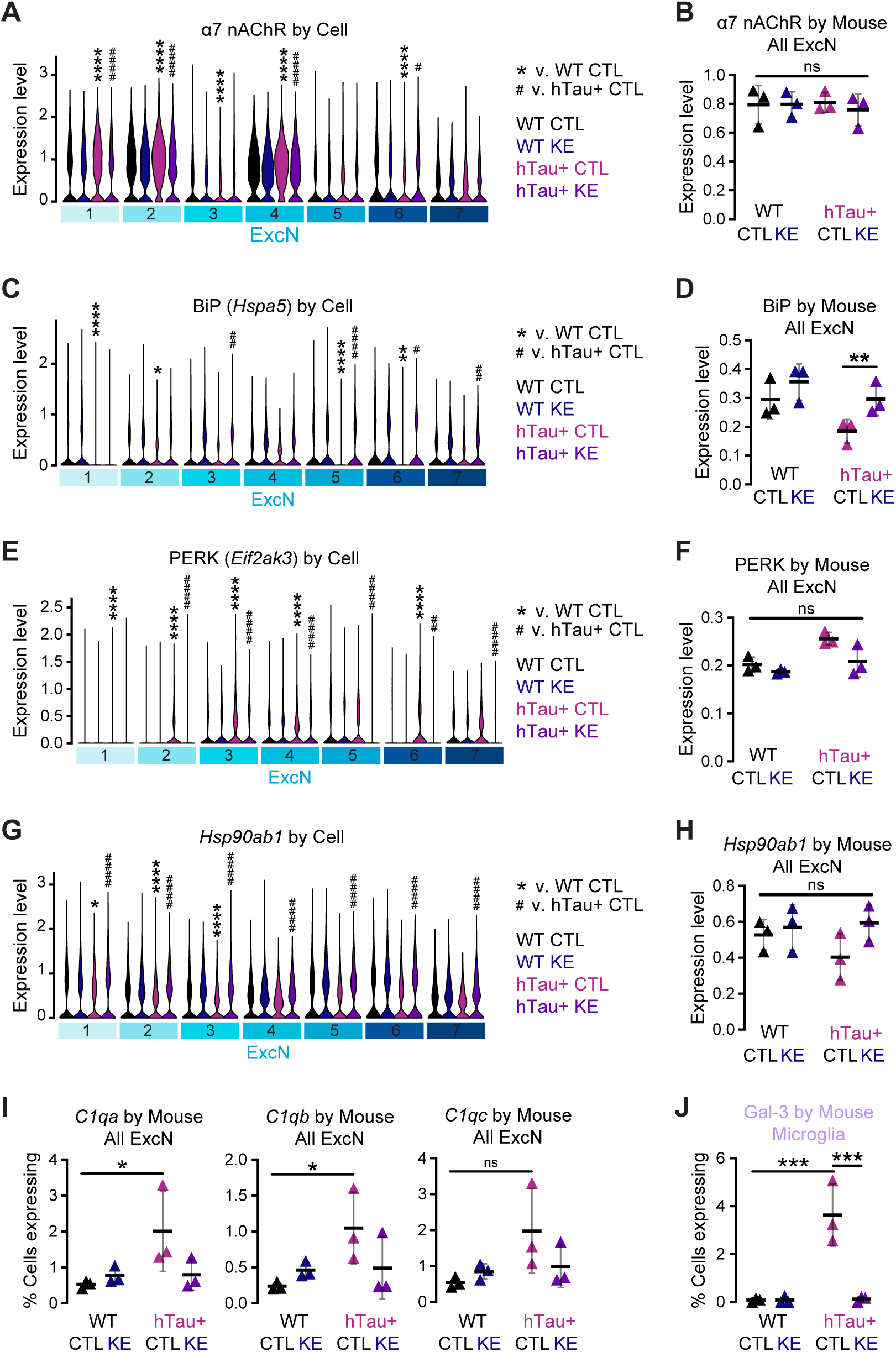
Analysis of ExcN clusters illustrates KE diet reversal to WT levels of synaptic and proteostatic status of ExcNs in hTau+ mice. **(A-H)** Relative expression levels of *α7 nAChR* (A-B), *Hspa5* (BiP) (C-D), *Eif2ak3* (PERK) (E-F), and *Hsp90ab1* (G-H) represented both at the single-cell level by ExcN cluster (A, C, E, G) and at the mouse level by pseudobulking (B, D, F, H). **(I)** Percentage of total ExcNs expressing synaptic markers *C1qa*, *C1qb*, and *C1qc*, quantified at the mouse level. **(J)** Percentage of total microglia expressing phagocytosis marker *Lgals3* (Gal-3), quantified at the mouse level. Data at the mouse level are represented as mean ± SD. * p < 0.05, ** p < 0.01, *** p < 0.001, **** p < 0.0001 relative to WT CTL, and # p < 0.05, ## p < 0.01, #### p < 0.0001 relative to hTau+ CTL, unless comparisons are specified. ns not significant. DESeq2 test was used for expression level comparisons (A-H) and Šídák’s multiple comparisons test for percent expression comparisons (I-J).

**Figure S5, related to Figure 5:**
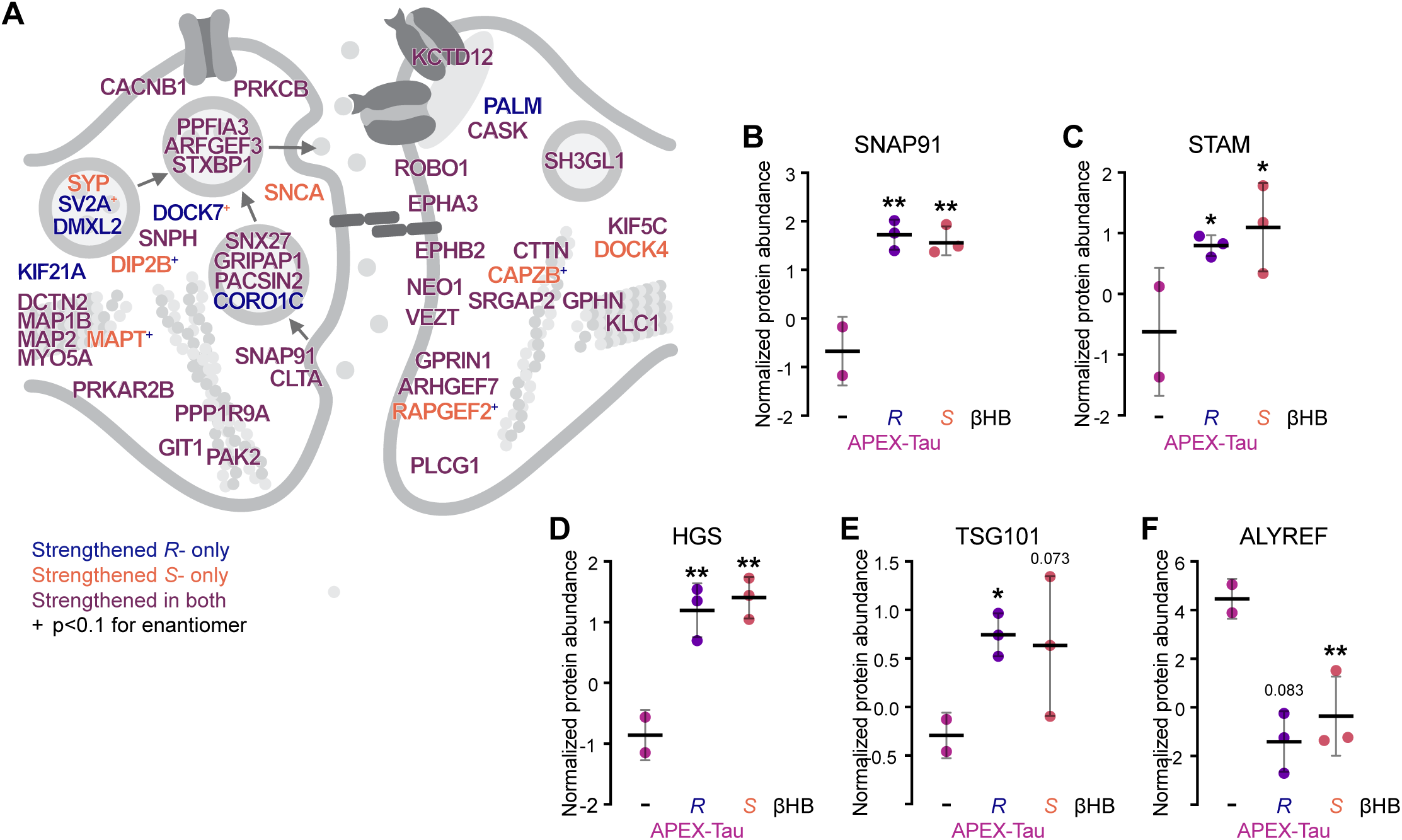
βHB alters neuronal Tau interactome, strengthening Tau’s interaction with endomembrane and synaptic systems. **(A)** Cartoon showing subcellular location of manually curated synaptic system proteins whose interactions with Tau are strengthened with βHB. The color of the protein signifies the enantiomer of βHB that is significant. Significance cut-offs were defined as p < 0.05 and |FC| > 0.5. The “+” indicates that the change in interaction was significant in one enantiomer of βHB while the other enantiomer had a p-value < 0.1. **(B-F)** Normalized protein abundances of Tau interactors SNAP91 (B), STAM (C), HGS (D), TSG101 (E), and ALYREF (F) with βHB treatments. P-values and FCs were determined by linear regression model to remove variance shared with APEX-Tub, representing artifactual interaction changes due to changes in the background proteome (see **Methods**). Data are represented as mean ± SD. * p < 0.05, ** p < 0.01, *** p < 0.001, **** p < 0.0001 by linear regression model.

**Figure S6:**
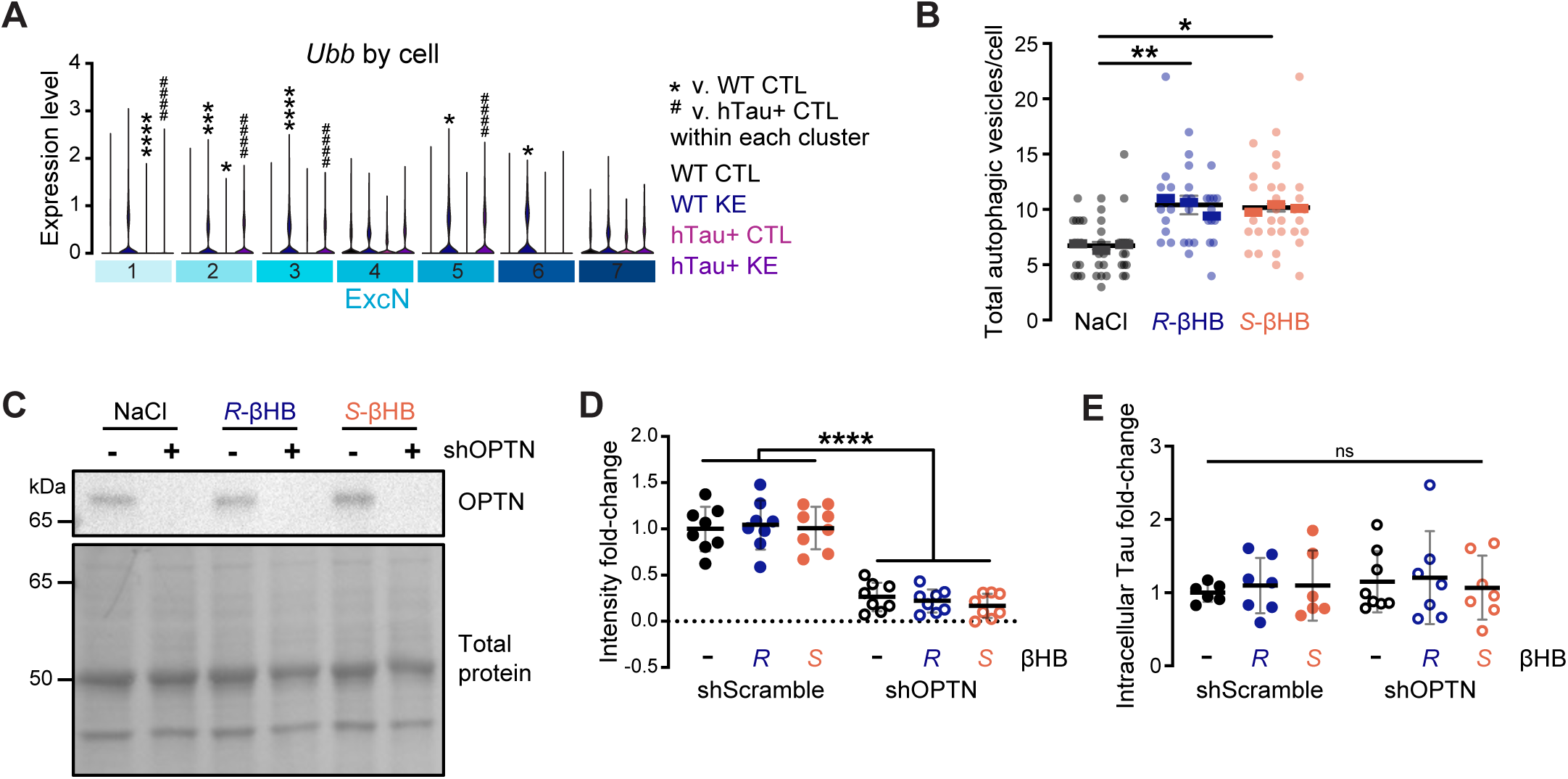
βHB enhances *Ubb* expression and increases autophagic vesicles in neurons; OPTN knockdown in neurons. **(A)** Relative expression levels of *Ubb* at the single-cell level by ExcN cluster. * p < 0.05, *** p < 0.001, and **** p < 0.0001 relative to WT CTL, #### p < 0.0001 relative to hTau+ CTL. *Ubb*, ubiquitin. **(B)** Quantification of total autophagic vesicles in primary neurons, corresponding to Fig. 6F**-H** (autophagosomes + autolysosomes). Each point represents an individual Map2+ neuron, with cells in the same well stacked into one column. Thick, color-coded bars represent the well mean (n = 8-15 cells/well), and black bars represent the overall group mean ± SD (n = 3 wells/group, from separate batches). * p < 0.05, ** p < 0.01 by linear mixed-effects model with Tukey *post-hoc* test. **(C-D)** Representative Western blot (B) and quantification (C) of OPTN in primary neurons, corresponding to Fig. 6I. Each point represents one independent well, normalized to the average of the NaCl-treated values for its respective plate. **** p < 0.0001 by Welch’s two-way ANOVA. **(E)** Quantification of relative intracellular Tau levels in primary neurons, corresponding to Fig. 6I. ns not significant by Šídák’s multiple comparisons test. Data for D-E are represented as mean ± SD.

## METHOD DETAILS

### Mouse genetics and husbandry

All animal procedures were carried out under guidelines approved by the Institutional Animal Care and Use Committee of the University of California San Diego. Mice were housed in a pathogen-free facility with a 12-hr light/dark cycle and ad libitum access to food and water. Standard vivarium chow was PicoLab Rodent Diet 20 (LabDiet #5053). All mice in this study were socially housed. This study primarily employed the PS19 mouse model of tauopathy (B6;C3-Tg(Prnp-MAPT*P301S)PS19Vle/J, Jackson Labs #008169), which expresses human mutant Tau^1N4R-P301S^ under the direction of the mouse *Prnp* promoter, as previously established.^47^ The mouse line was maintained by breeding hemizygous carrier males with non-carrier females on the original mixed C57BL/6J x C3H background. Genomic DNA was isolated using the AccuStart II PCR Genotyping Kit and amplified by PCR with AccuStart II GelTrack PCR SuperMix (Quantabio #95135, 95136). Primers used for PS19 genotyping are Forward (WT and hTau+): 5′ TTG AAG TTG GGT TAT CAA TTT GG 3′, Reverse (WT): 5′ TTC TTG GAA CAC AAA CCA TTT C 3′, Reverse (hTau+): 5′ AAA TTC CTC AGC AAC TGT GGT 3′. Post-mortem, tail snips were collected to confirm genotype. Only male mice were used in the current study, as they develop earlier and more consistent Tau pathology than PS19 females. Experimental takedowns were all done between 12-4 PM, unless otherwise specified, which is Zeitgeber Times (ZT) 6-10, with ZT0 representing lights on and ZT12 lights off.

### Diet studies

#### Ketone ester diet paradigms

The present study employs a precursor strategy for long-term supplementation of mice with βHB in lieu of supplementing βHB directly to avoid chronic high sodium load (e.g., by NaβHB) and provide a low-grade, sustained ketosis (**Fig. S1A**). The present study utilizes the ketone ester bis hexanoyl *R*-1,3-butanediol, abbreviated as C6×2-*R*-1,3-BD or C6×2-BD and referred to herein as “KE”.^48^ Stoichiometrically, one C6×2-BD produces 4 *R*-βHB: C6×2-BD is first hydrolyzed to two six-carbon medium chain fatty acids (FAs) and one *R*-1,3-BD. The *R*-1,3-BD is oxidized to one *R*-βHB in the liver by alcohol and aldehyde dehydrogenases, and the FAs undergo beta oxidation and ketogenesis in the liver to produce three additional *R*-βHB. Metabolism of C6 FAs to generate βHB occurs regardless of fed state, dietary carbohydrate content, or insulin status. βHB precursor diets are not reported to have any major side-effects, and we note none of concern in the present study.

Custom diets were designed and manufactured by Teklad. The KE diet (TD.200191) is 20% C6×2-BD by weight, substituting for carbohydrate content. Protein intake, non-KE fat intake, fat sources, micronutrients, fiber content, and other variables would be similar at equal caloric consumption to the control diet (“CTL”) TD.150345. C6×2-BD contains approximately 8 kCal/g. The diets were not designed to be equal in caloric density (kcal per gram of diet), as this would require using large amounts of non-caloric filler that inevitably have biological activity (e.g., fiber). Instead, the dietary components other than carbohydrates and KE were normalized on a per-calorie basis, and food intake was monitored closely. To assess bioavailability of βHB on the KE diet, 8 week-old male C57BL/6J mice were randomly assigned to CTL or KE diets and circadian rhythm boxes set to one of four ZT: ZT0, ZT6, ZT12, and ZT18. Mice were acclimated to their diets and new circadian rhythms for four weeks, during which body weight (BW) and food consumption were monitored. One mouse died prior to study endpoint for unknown reasons, likely fighting, and was not included in analysis. At the four-week timepoint, blood was collected in EDTA-coated tubes, and plasma was isolated by centrifugation then flash-frozen. At a later time, plasma *R*-βHB was measured using Cayman Chemicals βHB assay (#700190) following manufacturer’s protocol, which uses BDH1 enzyme to metabolize βHB to AcAc then measures the reaction byproduct, NADH.

For the therapeutic diet study in **Fig. 1-3**, PS19 transgenic (“hTau+”) and non-transgenic littermate wild-type mice (“WT”) were randomized by cage to diet conditions and switched from standard vivarium chow to either KE or CTL chow between 6-6.5 mo, when Tau pathology first presents in this model. Mice remained on the study diets for 15 weeks, until approximately 9.5-10 mo, at an advanced but pre-mortem disease state. Mice were monitored weekly for BW and food consumption (averaged per mouse by cage), and mice losing more than 25% of their initial body weight beyond week 1 were euthanized and excluded from additional analyses (n = 1 hTau+ CTL, n = 1 hTau+ KE, both dropped >25% of initial BW only at the endpoint weight measurement). Mice were also monitored for signs of severe pain or distress, including severe hindlimb paralysis that would prevent normal feeding, and only one mouse (WT CTL) was euthanized prior to study endpoint for fighting and excluded from all analyses. No mice died naturally prior to study completion. Post-mortem blood was collected from a separate set of WT and hTau+ mice on the same diet paradigm at ZT18.

#### Tissue preparation

At study endpoints, mice were anesthetized with a ketamine-xylazine cocktail, and toe and tail pinch reflexes were used to assess if the mouse was anesthetized deeply enough to proceed with perfusion. When the mouse was unresponsive to a pinch, the mouse was transcardially perfused with PBS for >3 min and until the flow-through was clear. After perfusion, mouse brains were extracted on ice. For diet study, one hemi-brain was fixed with 4% paraformaldehyde (PFA) for 36-48 hr, and the other hemi-brain was microdissected, with the cortex and hippocampus then flash-frozen and stored at −80°C for future biochemical analysis. After fixation, the brains were placed in 30% sucrose until each sank to the bottom of the container. On a sliding microtome, the mouse brain was mounted on OCT, and 30 μm-thick coronal sections were collected from the entire hippocampus, rostral to caudal. The sections were suspended in cryoprotectant, prepared with 30% sucrose, 1% PVP-40, and 30% ethylene glycol in 0.1 M phosphate buffer. The sections were stored in −20°C until further use.

### Biochemical experiments

#### Sequential protein fractionation

Mouse cortical tissue underwent homogenization and centrifugation in increasingly insoluble buffers - RAB, RIPA, then formic acid (FA) - to fractionate Tau species. RAB buffer stock was prepared as 0.1 M MES, 0.75 M NaCl, 1 mM EGTA, and 0.5 mM MgSO_4_, and inhibitors - 40 μM Na_3_VO_4_, 20mM NaF, 1:100 dilution protease inhibitor cocktail (PI; Sigma, #P8340), 1:100 dilution phosphatase inhibitor cocktail 2 (PIC2; Sigma, #P5726), 1:100 dilution phosphatase inhibitor cocktail 3 (PIC3; Sigma, #P0044), 1 mM PMSF, and 1.6 nM Trichostatin A (TSA; Sigma, #T1952) as an HDAC inhibitor - were added fresh. RIPA buffer stock was prepared as 50 mM Tris-HCl pH 7.4, 150 mM NaCl, 1 mM EGTA, 1 mM EDTA, 1% TritonX-100, 0.1% SDS, and 0.25% sodium deoxycholate, and inhibitors - 1:100 dilution PI, 1:100 dilution PIC2, 1:100 dilution PIC3, 1 mM PMSF, and 1.6 nM TSA - were added fresh. First, frozen tissue was combined with 10 μL of RAB buffer per mg of tissue, homogenized with a handheld bio-vortexer homogenizer, then sonicated until no chunks remained. Lysate was then centrifuged at 50,000x*g* at 4°C for 20 min, and the supernatant “RAB-soluble fraction” was frozen at −80°C. The pellet was combined with 10 μL of RAB buffer per mg of original tissue, subjected to the same extraction protocol, and the supernatant “RIPA-soluble fraction” was frozen at −80°C. Finally, the pellet was combined with 70% formic acid (FA) using 5 μL per mg of original tissue and subjected to the same extraction protocol. The FA-soluble supernatant was then neutralized 19:1 with neutralization buffer (1 M Tris-base, 0.5 M Na_2_HPO_4_), concentrated with Amicon Ultra Centrifugal Filters (MilliPore, #UFC510096) following manufacturer’s protocol, and the final “FA-soluble fraction” was frozen at −80°C.

#### Western blotting

To probe for proteins other than Tau (Tau protocol described in “*Sequential protein fractionation*”), cortical and cell lysates underwent homogenization and sonication in RIPA buffer directly, containing 1:100 dilution PI, 1:100 dilution PIC2, 1:100 dilution PIC3, 1 mM PMSF, 0.5 mM NaF, and 1.6 nM TSA. (Product codes for inhibitors in “*Sequential protein fractionation*.”) Lysates were centrifuged at 16,000x*g* at 4°C for 15 min, and supernatants were collected. Protein concentrations were determined by the BCA assay (Thermo, #23225). Equal amounts of protein were resolved on 4-12% Bis-Tris gels (ThermoFisher, NuPAGE system), transferred to either nitrocellulose or PVDF membranes depending on application, and probed with the specified antibodies. Total protein was visualized by Ponceau S or Imperial (LifeTech, #24615) stain. Bands in the immunoblots were detected using an enhanced chemiluminescence kit (Thermo, #34094) and visualized using a ChemiDocMP Imaging System (Bio-Rad). Band intensity was quantified using ImageJ. Representative blots from the same membrane are shown. Antibodies and dilution used as follows: DAKO (kindly provided by Peter Davies, The Feinstein Institute for Medical Research; 1:20000), PHF1 (Peter Davies; 1:1000), AT8 (Invitrogen, #MN1020; 1:1000), HT7 (Invitrogen, #MN1000; 1:1000), Tau5 (Invitrogen, #AHB0042; 1:1000), OPTN (ProteinTech, #10837-1-AP; 1:1000), LC3B (Millipore, #L7543; 1:1000), actin (Millipore, #A2066; 1:2000), GAPDH (CST, #2118; 1:2000), ubiquitin (SCBT, #sc-8017; 1:200), and goat HRP-conjugated secondaries (Invitrogen, anti-Rb: #31460, anti-Ms: #31430).

#### Immunofluorescence

The mouse brain slices were put into 24-well plates and gently washed with cold, sterile 1xPBS 3 x 5 min. The tissues were blocked with 10% normal goat serum, shaking for 1 hr at room temperature, then incubated overnight at 4°C in primary antibodies diluted in 0.4% PBS-Triton X-100 (PBST). The next day, tissues were taken to room temperature and left shaking for 30 min to allow binding of any leftover antibody. The secondary antibodies were also diluted in 0.4% PBST and left on the tissues shaking at room temperature for 1-2 hr. Finally, the tissues were washed in PBS containing DAPI stain 1×10 min followed by 2×5 min PBS washes. The stained tissues were then mounted and coverslipped. Antibodies and dilution used as follows: MC1 (Peter Davies, 1:1000), Iba1 (FujiFilm, #019-19741, 1:1000), and GFAP (Invitrogen, #PA1-10004, 1:1000).

Images were acquired on a Keyence BZX-700 fluorescent microscope with consistent settings within a single experiment. Mice and images with exclusively vasculature staining, indicating poor perfusion, were excluded from image analysis, but were included in biochemical analysis. All images within a single experiment were processed the exact same way in ImageJ, and the researcher conducting the analysis was blinded to the image identities. Minor brightness and/or contrast adjustments were performed as necessary, consistently across all images in the same experiment. As much as possible, auto thresholding was used to define the signal to be analyzed (most commonly using the max entropy algorithm), but when auto thresholding did not accurately capture the signal, manual thresholding was also used, including to exclude artifacts. Following thresholding, percent area and puncta analyses were performed. For graphing and statistical analyses, values were normalized to the WT CTL group to show a relative comparison across the four groups, unless for an hTau-specific comparison, in which case values were relative to hTau+ CTL group.

#### TauRD biosensor assay

Mouse cortical tissue was homogenized and suspended in PBS to prepare lysate for treatment in the TauRD biosensor assay.^52^ The following inhibitors were added to the PBS: 1:100 dilution PI, 1:100 dilution PIC2, 1:100 dilution PIC3, 1 mM PMSF, 20 mM NaF, 1.6 nM TSA, and 0.02% NaN_3_. (See inhibitor product codes in “*Sequential protein fractionation.*”) Frozen tissue was combined with 10 μL of PBS buffer per mg of tissue, homogenized with a handheld bio-vortexer homogenizer, then centrifuged at 10,000x*g* at 4°C for 10 min. The supernatant “PBS-soluble fraction” was frozen at −80°C until further use. Human TauRD P301S FRET biosensor-expressing HEK293T cells (ATCC CRL3275) were maintained and passed in complete DMEM media (DMEM with 10% FBS, 100 μg P/S). The cells were plated at a density of 125,000 cells/well in 500 μL of media on PLL-coated 10 mm coverslips in a 24-well plate. After 16 hr, the media was replaced with OptiMEM, and the cells were treated with lysate. Each lysate treatment was prepared from the PBS fraction at a concentration of 1 μg/μL through Pierce BCA Protein Assay Kits and diluted with 1x PBS buffer. For transfection, 25 μL of OptiMEM with 1.5 μL P3000 reagent (Thermo, #L3000008) was added to 7.5 μL of lysate and incubated for 10 min. Next, 25 μL of OptiMEM with 1 μL of Lipofectamine 3000 reagent (Thermo, #L3000008) was added to the lysate, and the total volume of 60 μL was added to each respective well dropwise. Post-24 hr treatment, the media was replaced with complete DMEM. After 48 hr or until the cells reached 85% confluency, the media was removed, and the cells were fixed with 4% PFA and stained with DAPI. The coverslips were mounted onto slides, and images were taken using Olympus VS200 Slide Scanner. 4 images of each coverslip were analyzed in ImageJ for quantification.

#### Protein overexpression and immunoprecipitation in HEK293T cells

pOPTN-EGFP was a gift from Beatrice Yue (Addgene, #27052).^97^ mScarlet-Tau constructs were made by cloning either mScarlet-Tau^WT^ or mScarlett-Tau^V337M^ into a pN1 plasmid backbone. HEK293T cells were maintained in complete DMEM media (DMEM with 10% FBS, 100 μg P/S) and used at a low passage number. At 60-70% confluency, cells were transfected with the DNA constructs using Lipofectamine 3000 reagents (Thermo, #L3000008), following manufacturer’s protocol. 1 hr before transfection, media was replaced with OptiMEM and then replaced with complete DMEM media 4 hr after transfection. After 24 hr, protein expression was checked with a fluorescent microscope, and experiments were performed when cells were at least 85% confluent and had strong expression of desired constructs.

For the Ub-Tau immunoprecipitation (IP) experiment, HEK293T cells were transfected with mScarlett-Tau^V337M^ and, for the Tau-OPTN co-IP experiment, HEK293T cells were co-transfected with OPTN-eGFP and either mScarlet-Tau^WT^ or mScarlett-Tau^V337M^. In both paradigms, cells were treated with Na*R*-βHB (Sigma, #298360), Na*S*-βHB (Cayman Chemicals, #21822), or NaCl as a control for 1 hr. Cells were lysed on ice in a nondenaturing lysis buffer: 50 mM Tris HCl pH 7.5, 150 mM NaCl, 150 mM MgCl_2_, 1% (v/v) NP-40, and 0.5% (w/v) sodium deoxycholate in milliQ water with inhibitors described in “*Western blotting*” added. Protein concentrations were determined by the BCA assay, and 150 μg of each sample was diluted to equal volumes of 500 μL in lysis buffer. To each sample, 1.5 μg of HT7 antibody (Invitrogen, #MN1000) was added, and samples mixed by rotation overnight at 4°C. Dynabeads G (Invitrogen, #10003D) were equilibrated to the lysis buffer, then 180 μg was added to each sample and mixed by rotation for 1 hr. Beads were pelleted by magnet, washed 3x with lysis buffer, then boiled at 95°C in 1x LDS and reducing buffer (Invitrogen, #B0007 and #NP0009) for 10 min. The supernatant is the IP sample, and input samples were prepared at equal concentrations. Immunoblotting was performed as described in “*Western blotting*.”

### Single-nuclei RNA sequencing

#### Nuclei preparation & sequencing

For single nuclei (sn)RNA sequencing analysis, three mice from each diet group (same mice as **Fig. 1**) were assessed. Hippocampi were isolated as described in “*Tissue preparation*” and stored at −80°C until further processing. Frozen tissues were dounce-homogenized in 3 mL lysis solution - 0.25 M sucrose, 25 mM KCl, 5 mM MgCl_2_, 10 mM TrisCl (pH 8), 10% Triton, and 0.1 M DTT with Protector RNase Inhibitor (Roche, #03335402001) - 40 times with the loose pestle and 40 times with the tight pestle. Then, nuclei isolation was performed with an iodixanol (Sigma, #D1556) gradient. Briefly, the lysed suspensions were mixed with 2 mL of 50% iodixanol, then added to the top of the iodixanol gradient (2.5 mL 25% iodixanol, 2 mL 30% iodixanol, 2 mL 40% iodixanol). The diluent for each iodixanol layer was 0.25 M sucrose, 25 mM KCl, 5 mM MgCl_2_, and 10 mM TrisCl, except for 50% iodixanol, which was diluted with 150 mM KCl, 30 mM MgCl_2_, and 120 mM TrisCl. The 30% layer also contained 1% phenol red. The gradient was centrifuged at 3184x*g* for 30 min at 4°C (acceleration 6, deceleration 1). To remove the nuclear layer, 1000 μL was removed from between the 30% and 40% iodixanol layers with a P1000 and collected in a 5 mL LO-BIND tube. 10 μL of nuclei suspension was used to count nuclei using Trypan blue. Then, the nuclei were resuspended in 3 mL of wash buffer (0.25 M sucrose, 25 mM KCl, 5 mM MgCl_2_, 10 mM TrisCl, 0.1% Tween-80) and centrifuged at 500x*g* for 10 min at 4°C. Supernatant was removed, and nuclei were resuspended once more in wash buffer with 1% BSA to achieve a concentration of 1200 nuclei/μL. Approximately 20,000 nuclei/sample were used for GEM generation, barcoding, and cDNA libraries using the Chromium GEM-X Single Cell 3’ Kit (ver. 4) and corresponding protocol. Pooled libraries were sequenced using the NovaSeq X Plus 25B in order to achieve >20,000 reads/nuclei.

#### Data preprocessing

A custom reference genome was constructed using CellRanger mkref (ver. 9.0.1) to add the hTau^1N4R-P301S^ transgene into the GRCm39 (2024-A) genome assembly as *hTau*.^98^ Fastq files were aligned to our custom PS19 genome using the CellRanger Count (ver. 9.0.1) pipeline using default settings and including introns.^98^ All samples passed basic QC at this stage. CellRanger output H5 files for each mouse were then preprocessed separately with the following steps: 1. Removal of ambient mRNA, 2. Filtering for healthy nuclei, 3. Removing homotypic doublets. First, cell-free mRNA contamination was estimated using the SoupX automated pipeline (ver. 1.6.2), which estimates the amount of ambient mRNA in droplet-based sequencing by profiling the empty droplets, compares it to droplets with nuclei, and adjusts mRNA counts accordingly.^99^ The SoupX output was then made into a Seurat (ver. 5.3.0) object for further processing.^100,101^

Second, healthy nuclei were filtered to remove nuclei where less than 200 genes, greater than 6,500 genes, or greater than 30,000 reads were detected. Genes that were expressed in fewer than 3 nuclei were filtered out. Moreover, any nuclei expressing more than 5% mitochondrial gene counts or more than 5% ribosomal gene counts were also removed. Data were log-normalized using the NormalizeData() function and the top 2,000 variable genes were identified using the FindVariableFeatures() function with the vst selection method. All genes were then scaled to unit variance and zero mean using the ScaleData () function. Principal component analysis (PCA) was performed on the variable feature set, and the first 15 principal components (PCs) were used to construct a shared nearest neighbor (SNN) graph. Clustering was performed using FindNeighbors() and FindClusters() (resolution = 0.04). Uniform Manifold Approximation and Projection (UMAP) was used for dimensionality reduction and visualization.

Third, multiplets of identical cell types, or homotypic doublets, were removed with DoubletFinder (ver. 2.0.6) following the standard pipeline.^102^ pN, or the number of generated artificial doublets, was left at the default 0.25. For 20,000 nuclei, the estimated doublet rate is 8%, and this was used to calculate the estimated total number of doublets, nExp. pK, or the principal component neighborhood size, was adjusted for each object using the paramSweep function and choosing the pK value with the highest bimodality coefficient. Finally, Seurat objects were merged and the layers joined with the JoinLayers function.

Heterotypic doublets were removed manually with a conservative strategy following the logic that any cell that highly expresses markers from two distinct cell types is likely a heterotypic doublet. Based on preliminary clustering and *a priori* knowledge of cell types in the hippocampus, we expected all cells to belong to one of the following classifications: neurons, oligodendrocytes, astrocytes, microglia, OPCs, and vasculature (e.g., endothelial cells and pericytes). We used our preliminary clustering to estimate the distribution of cell types in our study, which aligns with previously published studies and accepted estimates: 62% neurons, 18% oligodendrocytes, 9% astrocytes, 6% microglia, 3% OPCs, and 1.5% vasculature.^103–105^ We assigned a score for each of these six classifications to every cell based on cell-type specific markers highly expressed in our dataset. We used the assumption that cells of a particular cell-type would be the highest expressers of these genes and allowed for 5% leeway, e.g., if we expected 60% of the cells to be neurons, all neurons should be greater than the 60th percentile for neuron score, so we set our neuron score cutoff to the 65th percentile of neuron scores. Using this strategy, we removed 17.5% of predicted heterotypic doublets and 2% of cells with no score. This strategy did not significantly differentially identify and remove “doublets” from any particular group or cell type. As another validation of this strategy, we performed differential gene expression analysis between cells labeled using a DESeq2 test with FindMarkers and found that the differentially expressed genes (DEGs) between “singlet” and “doublet” within each group and cell cluster were almost exclusively cell-type specific markers for cell types not assigned to that particular cluster.

Lastly, we performed final clustering using 15 PCs and a resolution of 0.08 to achieve 14 clusters. Cell clusters were annotated by expression of known cell-type specific marker genes. For each cluster, the FindMarkers() function was used to identify differentially expressed genes. Enriched marker genes for each cluster were examined, and cluster identity was assigned by comparing marker gene profiles to canonical cell type markers reported in the literature. Cluster identities were assigned based on the following marker genes: microglia (*Cx3cr1*, *Csf1r*, *Fyb*), astrocytes (*Aqp4*, *Apoe*, *Slco1b2*), oligodendrocytes (*Mbp*, *Mog*, *Mobp*), OPCs (*Pdgfra*, *Neu4*, *Epn2*), vasculature (*Vtn*, *Vim*, *Pdgfrb*, *Pecam1*, *Emcn*, *Cldn5*), inhibitory neurons (*Gad1*, *Gad2*, *Reln*), excitatory neurons (*Slc17a7*), and DG neurons (*Prox1*, *Calb1*).

Though not explicitly mentioned in these methods, the packages Seurat.Utils (ver. 2.8.5) and scCustomize (ver. 3.0.1) were also used in pre- and post-processing.

#### Data analysis

Differential gene expression analysis was performed within each cluster at two levels, cell and mouse, and the same three comparisons were made at each level: WT CTL v. hTau+ CTL, WT CTL v. WT KE, and hTau+ CTL v. hTau+ KE. At the cell level, DEGs were determined using a DESeq2 test with FindMarkers. Significant DEGs were any with an adjusted p-value < 0.05 and were expressed in >10% of cells in at least one of the two groups in the comparison. At the mouse level, counts within a single cluster were pseudobulked, i.e., each gene was averaged within a cluster to the mouse level, using the AggregateExpression function, and then a DESeq2 test was performed. Significant DEGs were any with an adjusted p-value < 0.05. A separate analysis was also done with all seven ExcN clusters pseudobulked together with the same protocol. DEGs that overlapped in ≥4 ExcN clusters were further prioritized for analysis. Gene ontology (GO) analysis of this subset of DEGs was done in R using enrichGO in the clusterProfiler package (ver. 4.12.6), and the clusterProfiler simplify method was used to reduce redundancy of enriched GO terms.

Cellular metabolic states were modeled using Compass, a flux balance analysis algorithm developed by Wagner, et al.^56^ The Compass algorithm takes an input of single-cell transcriptomic profiles and a metabolic network (in this study, Mouse1^106^) and assigns a score by cell for each metabolic reaction based on the cell’s ability to maintain the reaction in the positive (and negative, if bidirectional) direction, i.e., flux. Compass outputs a list of reactions by cell with penalty scores, the higher the score meaning the less predicted metabolic flux through that pathway. These scores are then -log transformed, and the final values are “reaction scores,” with scores closer to zero representing greater metabolic flux. Compass was performed on ExcN clusters only in Module-Compass mode with the following subsystems specified: butanoate metabolism, sugar metabolism (glycolysis/gluconeogenesis, fructose and mannose, galactose, starch and sucrose, amino sugar and nucleotide sugar, and N-glycan), and lipid metabolism (all fatty acid terms, all beta-oxidation terms, all glyceride terms, and carnitine shuttle). To reduce scarcity and maximize capturing within-animal cell-to-cell variability, cells within an ExcN cluster and mouse were “micropooled” by random sampling of 100 cells without replacement, and counts were pseudobulked. For the butanoate subsystem, binary comparisons of reaction scores were performed comparing hTau+ CTL to WT CTL or to hTau+ by Welch’s t-test with a Bonferroni p-value correction. To aid interpretation of flux changes in bidirectional pathways, net reaction scores were calculated by subtracting the negative reaction score from the positive reaction score. Here, a positive net score represents net flux in the defined positive direction, and for metabolites net production, with larger score representing greater flux in the positive direction. A negative net score represents net flux in the defined negative direction, and for metabolites net consumption, with smaller (more negative) score representing greater flux in the negative direction. PCA in Compass space was performed with prcomp (stats, ver.4.4.3).

### Fly experiments

#### Fly husbandry and diets

Fly lines (*elav^C155^*–*Gal4*, *UAS-hTau^1.13^*) used in this study were obtained from the Bloomington Drosophila Stock Center (BDSC). Flies were maintained on standard diet (Nutri-Fly, #66-113, Genesee Scientific Corporation) in the lab for several generations. The *C155>UAS-hTau^1.13^* flies were generated by crossing *elav^C155^*–*Gal4* with *UAS-hTau^1.13^*. The βHB diets were made by supplementing the standard diet with 2 mM sodium βHB (*R*- or *S*-). The control diet was made by supplementing the standard diet with 2 mM NaCl.

#### Life span assay

*C155>UAS-hTau^1.13^* flies were collected after setting the cross. Flies were separated by sex, genotype, and diet. For each condition/group, 20 flies were collected into one vial, and there were 5 vials for each condition, which makes a total of 100 flies in each group. Day 0 was defined by the day that new adult flies were hatched. New flies were transferred to different diets on Day 1-2. The number of deaths in each vial was checked every two days.

#### Climbing assay

60 flies from same sex, genotype, and diet were collected and separated into 4 vials. All the flies were maintained on the desired diet starting from Day 1 after they hatched. Climbing assays were performed every 7 days starting from Day 7. Flies were transferred to an empty vial with standard diet at the bottom of the vial at room temperature. The measurement vial was divided into 6 equal regions representing climbing score. Each group of flies has a 15-sec testing period starting from tapping them down to the surface of the food. At the end of the test period, the scores for each fly are recorded and averaged. Flies were allowed to rest for 1 min. 3 technical replicates were performed, and the score from this group was determined by the average score of 3 technical replicates.

#### Brain slice culture experiments

Organotypic brain slice cultures (BSC) were performed with hippocampal slices obtained from wild-type P8-10 mouse pups, as previously described, with modifications.^107^ Pups were culled by decapitation, and bilateral hippocampi were quickly dissected in aCSF, as described in “*Tau secretion assays*,” that had been oxygenated. 300 μm thick slices were cut using a tissue chopper and cultured in Millicell inserts (Millipore, #PICM0RG50) in 6-well plates, 4-6 slices/insert, in BSC media: 50% (v/v) Basal Medium Eagle (BME), 25% (v/v) Heat Inactivated Horse Serum, 1% (v/v) GlutaMAX (Thermo, #35050-061), 0.5% (v/v) P/S (Thermo, #15070-063), 0.033% (v/v) insulin, 45 mM *D*-Glucose and 25 mM HEPES in EBSS buffer and sterile-filtered (0.2 μm).

On DIV0, slices were transfected with AAV2-hTau^P301S^ (ViroTek) at a titer of 2E10 vg/mL. On DIV2, slices were treated with 1.5 μg/mL homemade K18-Tau^P301L^ pre-formed fibrils (PFF). On DIV4, Na*R*-βHB, Na*S*-βHB, or NaCl (all 10 mM) was added to the media and refreshed with each media change, every 2-3 days. On DIV14, slices were fixed in 4% PFA at room temperature for 2-3 hr. Following fixation, slices were permeabilized overnight in 1% Triton X-100 in PBS. Antigen retrieval was performed using a citrate buffer (pH 6.0) under low pressure in a pressure cooker for 15 min. Slices were then blocked in 20% BSA with 0.4% Triton X-100 in PBS for 2-3 hr at room temperature. Slices were incubated with primary antibodies in 0.4% Triton X-100 in PBS for 48 hr at 4°C. Afterward, slices were washed six times for 10 min each in PBS, followed by incubation with fluorophore-conjugated secondary antibodies and DAPI for 2 hr at room temperature. Finally, slices were washed an additional six times for 10 min in PBS, mounted together with their membrane inserts onto glass slides, and coverslipped using Fluoromount-G. Images were acquired on a Keyence BZX-700 fluorescent microscope.

### Primary neuron experiments

#### Culture preparation

Primary cortical neuronal culture was prepared using E18 Sprague Dawley rats (Charles River Labs, #001) following previously described protocols. Briefly, the cerebral cortices of the embryos were dissected on ice, meninges removed, and collected in cold dissection media containing DAP-5 and NBQX. After dissection, tissues were washed in dissection media 1-2 times and digested in papain for 20 min at 37°C. The digestion was quenched with low OVO and DNase, then the tissues were triturated and filtered through a 70 µm cell strainer. Cell solution was pelleted by centrifugation, gently resuspended in high glucose media to remove debris, and pelleted again by centrifugation. Lastly, the cell pellet was resuspended in neuronal media (NB, B27, P/S, GlutaMAX; Thermofisher #12349-015, #17504-044, #15070-063, #35050-061 respectively) and plated in neuronal media at 500K per mL on plates precoated with poly-*D-*lysine (PDL; Thermo, #A3890-401). Cells were used for experiments between day-*in-vitro* (DIV) 8-12.

#### In vitro βHB treatments

The following reagents were used for the *in vitro* metabolite experiments in this study: Na*R*-βHB (Sigma, #298360), Na*S*-βHB (Cayman Chemicals, #21822), NaCl, and AcAc (Sigma, #123-54-6). Stock bottles were stored at temperatures according to manufacturer’s guide, and additionally, Na*R*-βHB stock container was stored in an air-tight container with Drierite desiccant, as Na*R*-βHB is highly hygroscopic. Metabolites were prepared in milliQ water at 1 M, and cells were treated at 10 mM (1:100) unless otherwise specified. Working solutions were stored at 4°C for up to four weeks, then prepared freshly again, and we note no discernable changes in efficacy due to storage conditions. Metabolite treatments were added directly to cell culture media unless media change was essential to experimental design (e.g., secretion assay).

#### Tau secretion assays

Artificial cerebrospinal fluid (aCSF) was prepared (140 mM NaCl, 5 mM KCl, 2.5 mM CaCl_2_, 2 mM MgCl_2_, and 10 mM HEPES), with and without 10 mM *D*-glucose (Sigma, #G8270). Cells were gently washed once in aCSF without glucose, then treated as specified for 1 hr. Conditioned media was then collected and spun down at 8,000x*g* for 5 min to remove any cell debris. Cells were then rinsed on ice in PBS and lysed in RIPA buffer with inhibitors (same as RIPA in “*Sequential protein fractionation*”), sonicated, then spun down to remove membranes and debris, as described in “*Western blotting*” section.

#### Tau ELISAs

Tau ELISAs were done as previously described.^108^ Briefly, high-bind plates (Corning, #3925) were coated overnight with BT2 antibody (1.5 μg/mL; Thermo, #MN1010) for capture of murine Tau. Plates were washed, blocked in 3% BSA in DPBS with Ca^2+^/Mg^2+^ (Thermo, #14287-072) for 3 hr at room temp, then pre-incubated for 1 hr at room temp with a standard curve of Tau441 (rPeptide, #T-1001), 1:4 dilutions of conditioned media, and 1:250 dilutions of cell lysate. Samples were run in triplicate. Tau5-AP was added (homemade following manufacturer’s protocol; Tau5: Abcam, #AB80579, AP kit: Bio-Rad, #LNK012AP), and the plates were left shaking overnight at 4°C. The following day, plates were brought to room temp to shake for one additional hour, then thoroughly washed. Chemiluminescent reagent CDP-Star with Sapphire-II (Invitrogen, #T2214) was added, and the plates incubated in the dark for 30 min then were read using a chemiluminescent-compatible plate reader. The standard curve was graphed in GraphPad Prism using a second order polynomial nonlinear regression, and unknowns were interpolated. Outlier wells were excluded from analysis.

#### LC3 reporter assay

Lentivirus expressing mCherry-GFP-LC3 was prepared as previously described.^109^ Primary neurons were plated at a density of 600K per mL onto PDL-coated coverslips and were infected on DIV3 with the homemade lentivirus-mCherry-GFP-LC3 for >5 days. Mature neurons were treated with *R*-βHB, *S*-βHB, or NaCl for 1 hr, then fixed. The cells were then blocked in 10% BSA, stained for MAP2 (Sigma, #AB5622) overnight then DAPI for 10 min, and mounted. A Nikon AXR confocal microscope system was used for imaging. 488 nm and 568 nm lasers were used to excite the GFP and mCherry, respectively, and all images were taken with the same confocal settings, including identical aperture and laser intensities, Galvano scanner settings, and number of scans averaging. To ensure cells included in analysis were transfected, only cells with at least one punctum were imaged. Only MAP2+ cells were analyzed. mCherry was pseudocolored in ImageJ to magenta for colorblind accessibility. Magenta and white (co-localization of magenta and green signal) puncta were manually counted in each cell. A linear mixed model was used to analyze the data (described in “**Statistical analysis**”), with each point representing the puncta count for a single cell, and all points in a column representing a single, independent well.

#### shRNA knockdown experiments

The plasmid pLKO.1-puro expressing the shRNA scramble construct was a gift from David Bryant (Addgene plasmid #162011).^110^ The plasmid pLKO.1-puro expressing shOPTN was purchased from Sigma’s MISSION shRNA (#SHCLNG) catalog, target sequence 5’-CTGAAAGAGAACAATGACATT-3’ (TRCN0000177595). Lentivirus expressing either construct was prepared as previously described.^109^ Primary neurons were plated at a density of 600K per mL onto PDL-coated 12-well plates or coverslips and were infected on DIV3 with the homemade lentivirus for 5 days. shOPTN efficacy was validated via Western blot with anti-OPTN antibody (Proteintech, #10837-1-AP). For the Tau secretion assay, the exact protocol as described in “*Tau secretion assays*” was followed.

For the Halo-Tau assay, primary neurons were co-transfected with shOPTN or shScramble and HaloTag-Tau^P301L^ lentiviruses on DIV3.^63^ On DIV7, cells were treated with dox to induce HaloTag-Tau^P301L^ construct expression. On DIV8, 500 nM of HaloTag ligand tetramethylrhodamine (TMR; Promega, #G8281) was added to culture media for 2 hr. Cells were then washed once with aCSF then incubated with 5 μM HaloTag blocker (non-fluorescent ligand 1-chloro-6-(2-propoxyethoxy)hexane; Astatech, #F83341) for 1 hr. Next, cells were washed once with aCSF and co-treated with βHB and 50 nM LysoTracker Far-Red (Invitrogen, #L12492) in aCSF with glucose for 1 hr, after which the cells were fixed with PFA. The cells were then blocked in 10% BSA, stained for MAP2 (Sigma, #AB5622) overnight then DAPI for 10 min, and mounted. Eclipse Ti2-E spinning disk confocal microscope was used for imaging, and 561 nm and 640 nm lasers were used to excite the TMR ligand and LysoTracker, respectively. ImageJ was used for analysis. To control for viral transfection efficacy and variability in viral load, only cells with discrete, countable puncta (i.e., cells were not entirely green/overloaded with puncta) were included in analysis. Three initial ROIs were made: all Halo-Tau via manually thresholding the red channel, all lysosomes by manually thresholding the far-red channel, and single neurons by freehand selecting MAP2+ cells. Two additional ROIs were made from the combination of these ROIs, and percent area was calculated from these measurements: Halo-Tau AND a single neuron for all the Tau within the cell (denominator), and Halo-Tau AND lysosomes AND a single neuron for all the Tau within the cell also within the lysosome (numerator). A linear mixed model was used to analyze the data (described in “**Statistical analysis**”), with each point representing the puncta count for a single cell, and all points in a column representing a single, independent well.

### iPSC-Neuron experiments

#### iPSC lines & maintenance

This study employed three induced pluripotent stem cell (iPSC) lines, all previously described: APEX2-N-Tau^2N4R^, APEX2-Tubulin α-1B (Tub), and a WT iPSC line.^63,64,111^ Each APEX2 line originates from the same iPSC line, WTC11 (non-diseased genetic background, XY), which was then engineered to possess a Tet-ON 3G-controlled Neurogenin-2 cassette integrated into one allele of the AAVS1 locus for inducible neuronal differentiation (i^3^Ns).^111^ Integrated into the other AAVS1 allele, APEX-Tau iPSCs possess Tet-ON 3G-controlled expression of APEX2 conjugated to the N-terminus of wild-type human Tau^2N4R^, and APEX-Tub iPSCs possess APEX2 conjugated to the N-terminus of human Tubulin α 1B. The WT iPSC line was used in “*HaloTag pulse tracing*” and is also a non-diseased genetic background, XY. All iPSC lines were maintained routinely in Essential 8 media (E8; LifeTech, #A1517001) in a cell culture incubator with 5% CO_2_ at 37°C. Upon reaching confluency, iPSCs were passed by dissociating with StemPro Accutase (LifeTech, #A1110501) and replating onto Matrigel-precoated plates (Corning, #356231) with 10 μM ROCK inhibitor (RI; Y-27632, Cayman Chemicals, #10005583) for 24 hr.

#### iPSC-Neuron differentiation

Human i^3^Ns were differentiated with a simplified two-step protocol: pre-differentiation and maturation. For pre-differentiation, iPSCs were replated in Matrigel-precoated plates in pre-differentiation media: knockout DMEM/F-12 (LifeTech, #12660-012), 1:100 N2 supplement (LifeTech, #17502-048), 1:100 non-essential amino acids (NEAAs; LifeTech, #11140-050), 2 μg/mL dox (Sigma, #D9891), 10 ng/mL brain-derived neurotrophic factor (BDNF; Peprotech, 450-02), 10 ng/mL neurotrophin-3 (NT-3; Peprotech, #450-03), 1 μg/ml mouse laminin (LifeTech, #23017-015), and 10 μM RI. Media was replaced daily, and RI was removed after one day. iPSCs took three days to fully pre-differentiate (i.e., D-3 to D0). On D0, the pre-differentiated precursor cells were dissociated with accutase and re-plated into PDL-precoated dishes, plates, or coverslips according to experimental need in maturation media: 50% NB, 50% DMEM/F12 (LifeTech, #11320-033), 1:200 N2, 1:100 B27, 1:100 NEAAs, 2 μg/mL dox, 10 ng/mL BDNF, 10 ng/mL NT-3, 1 μg/ml laminin, and 1:200 GlutaMAX. One-third of the media was replaced weekly with fresh maturation medium thereafter, minus dox. i^3^Ns were used for experiments after D28. Halo-Tau pulse tracing The HaloTag-Tau^P301L^ tool is described completely in Xiao, et al. (2026).^63^ Mature i^3^Ns overexpressing HaloTag-Tau were labeled with 1 μM HaloTag ligand tetramethylrhodamine (TMR; Promega, #G8281) for 1 hr by directly adding ligand to maturation media. Cells were then washed twice with maturation media then incubated with 5 μM HaloTag blocker (non-fluorescent ligand 1-chloro-6-(2-propoxyethoxy)hexane; Astatech, #F83341) for 30 min. Lastly, media was replaced with fresh media containing 5 μM HaloTag blocker and 10 mM of NaCl, *R-*βHB, or *S-*βHB for 1 hr, at which point cells were lysed on ice. Samples for SDS-PAGE and Western blot analysis were prepared as described in “*Western blotting*.” In-gel fluorescence of the TMR ligand was detected using a ChemiDoc imager (Bio-Rad). 4R-Tau (CST, #30328), actin (CST, #4967), and vinculin (CST, #4650) antibodies were used for immunoblotting.

### Mass spectrometry-based neuronal Tau interactomics

#### APEX labeling

The experimental design for the APEX2 study consisted of two lines, APEX2-Tau and APEX2-Tub, each undergoing three treatments, NaCl, Na*R*-βHB, and Na*S*-βHB, in biological triplicate, for a total of six groups and 18 independent samples. APEX2 fusion iPSC lines were differentiated into i^3^Ns on 10-cm^2^ dishes following the previously described two-step protocol in “***iPSC-Neuron experiments***,” with a density of 5 M of pre-differentiated precursor cells replated into each final 10-cm^2^ dish. APEX labeling was done in three batches of six plates, one plate per group, adapted from previous protocol.^64,112^ Batches were staggered by 1.5 hr, and treatments within a batch were staggered by 5 min. At D27 of maturation, i^3^Ns were treated with 2 μg/mL dox to induce expression of APEX-conjugated bait proteins. After 24 hr, i^3^Ns were washed once gently with aCSF with glucose, and treated for 30 min with 500 μM biotin-phenol (Adipogen, #CDX-B0270-M100) and 10 mM of either NaCl, Na*R*-βHB, or Na*S*-βHB in aCSF with glucose. Interactors were then biotin-labeled for exactly 1 min by addition of 1 mM H_2_O_2_. The labeling was quenched and cells were washed three times in a freshly-made, cold quenching solution of 10 mM sodium ascorbate, 5 mM Trolox (Sigma, #238813), and 10 mM sodium azide in 1x PBS (LifeTech, #70011-044). Lysis buffer stock consisted of 50 mM Tris, 150 mM NaCl, 0.1% (wt/vol) SDS, 0.5% (wt/vol) sodium deoxycholate, and 2% (vol/vol) TritonX-100, pH 7.5. 1 mL of lysis buffer plus inhibitors (1:100 dilution PI, 1:100 dilution PIC2, 1:100 dilution PIC3, 1 mM PMSF, 0.5 mM NaF, and 1.6 nM TSA) and quenchers (10 mM sodium ascorbate, 5 mM Trolox, and 10 mM sodium azide) was added per dish. (See inhibitor product codes in “*Sequential protein fractionation.*”) Cells were scraped, collected, sonicated, then centrifuged at 16,000x*g* at 4°C for 15 min. Supernatant was collected and stored at −80°C until ready for the next step.

#### Streptavidin bead pulldown

Lysates were thawed on ice, and protein was quantified per sample using a Pierce 660 nm assay (Pierce, #22660). 750 μg from each lysate was diluted to equal volumes (∼0.5 μg/mL) in RIPA buffer. Pierce Streptavidin Magnetic Beads (Thermo, #88816) were equilibrated in RIPA buffer, combined with lysates at a 1:1 protein:bead ratio, and mixed overnight at 4°C in a tube rotator. The following day, beads were pelleted on a magnetic rack (Invitrogen, #12321D) and thoroughly washed with 1 mL of each of the following: twice with RIPA buffer, once with 1 M KCl, once with 0.1 M Na_2_CO_3_, and twice with 2 M urea, 50 mM ammonium bicarbonate.^112^ A small portion (<5%) of the beads were set aside for Western blot validation, and the remainder were frozen at −80°C until ready for the next step.

#### Peptide purification and isobaric labeling

To prepare samples for mass spectrometry (MS) analysis, protein was digested on-bead with 1 mAU LysC (Fujifilm, #121-05063) and 300 ng trypsin (Promega, #V5113) in 2 M Urea, 50 mM Tris buffer (pH 8) at 37°C overnight with shaking. 10 mM TCEP and 10 mM iodoacetamide were added after digestion and allowed to incubate at room temperature for 30 min in the dark. On-bead digests were acidified to 1% trifluoroacetic acid (TFA), desalted via StageTips made in-house, and labeled with 18-plex tandem mass tags (TMTPro, Thermo, lot #A52045) using an on-column labeling protocol.^112,113^ For on-column TMT labeling, StageTips were washed with 100 μl of methanol, 100 μl of 50% ACN, 0.1% TFA, and equilibrated with 100 μl 0.1% TFA twice. The digest was loaded by spinning at 2000x*g* until the entire digest passed through. Resin-bound peptides were washed twice with 75 μl of 0.1% FA. 1 μl of TMT reagent in 100% ACN was added to 100 μl of 50 mM phosphate buffer (pH 8) and then added to the resin, centrifuging at 200x*g* for 4 min, then 1000x*g* for 1 min. The phosphate buffer and excess TMTs were washed away with 75 µl 0.1% FA twice, and peptides were eluted with 50 µl 50% ACN, 0.1% FA and then 50 µl 50% ACN, 20 mM ammonium formate (pH 10). Peptide concentrations across all 18 samples were estimated by Nanodrop A280 and mixed at 1:1 ratio. Mixed TMT-labeled peptides were dried via vacuum centrifugation and resuspended in 0.1% FA, then step-fractionated by basic reverse-phase fractionation. Two 18-gauge needle punches of sulfonated divinylbenzene (SDB-RPS; Empore #98060402231EA) were packed into a StageTip and then washed with 100 µl of methanol, 100 μl of 50% ACN, 0.1% FA, and equilibrated with 100 μl 0.1% FA twice. Peptides were step-fractionated into 9 100 µl fractions (5%, 10%, 15%, 20%, 25%, and a final fraction of 75 µl 30% ACN, then 25 µl 45% ACN in 20 mM ammonium formate (pH 10) buffer). Each fraction was dried separately via vacuum centrifugation and resuspended in 9 µl 0.1% FA for subsequent LC-MS/MS analysis.

#### LC-MS/MS

Peptides were analyzed by LC-MS/MS using an Orbitrap Eclipse Tribrid mass spectrometer (ThermoFisher Scientific) coupled online to an Easy nLC 1200 (ThermoFisher Scientific). 4 of total 9 µl of each fraction was loaded onto a 75 mm i.d. PicoFrit column (New Objective, Woburn, MA) packed with 1.9 mm AQ-C18 material (Dr. Maisch, Germany) to a length of 35 cm. A 110-min gradient was utilized with HPLC solvent A as 2% ACN, 0.1% FA and solvent B 80% ACN, 0.1% FA. The gradient went from 10% B at 1 min to 50% B by 85 min, then to 72% B at 94 min, 90% B between 95 and 100 min, then 60% B for the remainder of the runtime. A precursor scan was first run from 375-1800 *m/z* at resolution 60,000, with an automatic gain control (AGC) target of 4e5 and a maximum injection time of 50 ms. For precursor selection, monoisotopic peak determination was set to “peptide” with the most abundant peak used for the isolation window center. Ions were selected within a charge state range of 2-6, minimal intensity of 20,000, and dynamic exclusion was set to 20 seconds. Data-dependent scans were collected at a top speed for a 2 second duty cycle. Precursors underwent high-collision dissociation fragmentation with a normalized collision energy of 32, and product ions were measured in the Orbitrap at a resolution of 50,000. The precursor isolation window (quadrupole) was set to 0.7 *m/z*, and the maximum MS2 injection time was 105 ms, targeting an AGC of 200%.

#### Protein quantification and analysis

Data were searched with Spectrum Mill (Agilent) using the Uniprot Human database (Dec 28, 2017), containing 264 common laboratory contaminants and 553 smORFs. A fixed modification of carbamidomethylation of cysteine and TMTpro-18plex labels, and variable modifications of N-terminal protein acetylation, oxidation of methionine, and pyroglutamic acid were searched. The enzyme specificity was set to trypsin/P with a maximum of three missed cleavages. The MS1 and MS2 mass tolerance were set to 20 ppm. Peptide FDRs were calculated to be less than 1% using a reverse, decoy database. Reporter ion corrections based on the TMT label lot were applied, and peptides were summarized into protein groups, excluding precursor ion isolation purities of <50%. Proteins were grouped without expanding subgroups and ratioed based on the mean of signal across all channels.

Proteins which were identified by fewer than 2 unique peptides, or were common laboratory contaminants or human keratins, were filtered out of the data set. To assess data quality, data were clustered by columns (1 - Pearson correlation, average linkage via Morpheus) prior to Pearson correlation similarity matrix.^114^ Two samples (1 APEX-Tau x NaCl, channel 127N; 1 APEX-Tub x Na*S*-βHB, channel 129N) were excluded from all following analyses due to Pearson’s correlation coefficient r < 0.2 with other two replicates in the same group. Data normalization was performed using the global Median-MAD method via ProTIGY.^115^ To facilitate direct comparison among the NaCl and Na*R*-βHB/*S*-βHB treatment APEX-Tau samples, we removed variance attributable to the APEX-Tub samples by applying a linear regression model via the lmfit and eBayes functions in the R limma package. The resulting residuals for each comparison were then used for downstream analyses.

The following significance cut-offs were used for binary comparisons within APEX-Tau to compare Na*R*-βHB v. NaCl and Na*S*-βHB v. NaCl: nominal p-value < 0.05 and |FC| > 0.5. Proteins with FCs > 0.5 were considered significantly strengthened in their interaction with Tau upon βHB treatment, and conversely proteins with FCs < −0.5 were considered significantly weakened in their interaction with Tau upon βHB treatment. Gene set enrichment analysis (GSEA) was performed within each binary comparison using clusterProfiler gseGO.^116,117^ Protein lists were ranked by a score: -logP*FC. The clusterProfiler simplify method was used to reduce redundancy of enriched GO terms.

## STATISTICAL ANALYSIS AND DATA REPRESENTATION

Mouse experiments included at least two independent cohorts of mice, with the exception that a single cohort of mice was used for the bioavailability study in **Fig. S1A-B**. Littermates were randomized to diet intervention. All *in vitro* experiments were done on at least two independent multi-well plates per experiment with the control group represented on each plate for batch normalization. Data was normalized to the control group for the experiment, and this is specified throughout as “fold-change” and “relative” quantifications.

Each point on a graph represents a biological replicate, and technical replicates were summed to the most meaningful biological level. For mouse experiments, each point represents one individual mouse, and any repeated measurements from the same mouse, e.g., immunofluorescent images, were averaged. For the snRNA-seq experiments, some data are presented at the cell level, which is specified when applicable. For *in vitro* experiments analyzed by Western blot or ELISA, each point represents a single well. For the TauRD biosensor assay, each point represents value from one animal by the average of two technical duplicate wells, as described in “*TauRD biosensor assay*.” For all other imaging experiments, each point represents a single cell or brain slice, all cells/slices within a single well are stacked in the same column, and the mean value for the well is depicted. Each data point for the APEX experiments represents one 10-cm^2^ dish of treated i^3^Ns, and the labeling was done in batches of six with one plate per group to minimize batch effects. Group means are shown, and all error bars represent +/-standard deviation (SD) of the mean. Graphs were made in GraphPad Prism (ver. 10) and R (ver. 4.4.3), and aesthetics were adjusted in Adobe Illustrator (2024).

Statistical analysis was primarily performed in Prism. p < 0.05, p < 0.01, p < 0.001, p < 0.0001, and not significant represented as *, **, ***, ****, and ns, respectively, and represent pairwise comparisons unless otherwise specified. Statistical tests and multiple comparisons were decided *a priori* and kept consistent throughout experimental designs. For experiments with just two groups, e.g., hTau+ CTL v. hTau+ KE, Welch’s t-test was performed. For single-variable experiments with 3+ groups, e.g., NaCl v. *R*-βHB v. *S*-βHB, ordinary one-way ANOVA (Welch’s) and Dunnett’s multiple comparison tests were performed. For two-variable experiments, e.g., (hTau+ v. WT) x (CTL v. KE), full-model two-way ANOVA and Šídák’s multiple comparisons tests were performed. Before performing a two-way ANOVA, we tested for homogeneity of variance with Levene’s test in R (car package, ver. 3.1). If this assumption was violated, we proceeded with Welch’s ANOVA, which is more robust for cases of unequal variance. A linear mixed model was used for cell and brain slice culture imaging experiments, in which replicates within the same well are not independent but are included in analysis to more accurately represent the within-well biological variability.^118^ This analysis was done in R (ver. 4.4.3) with the lme4 (ver. 1.1-37) and emmeans (ver. 1.11.2) packages. Area under the curve analysis was calculated in Prism with default settings, then areas and SDs were compared by one-way ANOVA. The following comparisons were decided *a priori* based on biologically meaningful comparisons and were performed in every analysis as applicable: (1) All diet study analyses: WT CTL v. WT KE, WT CTL v. hTau+ CTL, and hTau+ CTL v. hTau+ KE. (2) All one-variable βHB analyses: NaCl v. *R*-βHB and NaCl v. *S*-βHB. (3) All two-variable βHB analyses (with Variable1, Var1, being βHB treatment): NaCl Var2A v. *R*-βHB Var2A, NaCl Var2A v. *S*-βHB Var2A, NaCl Var2A v. NaCl Var2B, NaCl Var2B v. *R*-βHB Var2B, and NaCl Var2B v. *S*-βHB Var2B. The exception is when a linear mixed-effect model was performed, as the emmeans package automatically makes all possible comparisons.

Statistical methods for the snRNA-seq study are described in “***Single-nuclei RNA sequencing***.” Statistical methods for the neuronal interactomics study are described in “***Mass spectrometry-based neuronal Tau interactomics***.”

## RESOURCE AVAILABILITY

### Lead contact

Requests for further information and resources should be directed to and will be fulfilled by the lead contact, Xu Chen.

### Materials availability

This study did not generate new unique reagents.

### Data and code availability

- The raw sequencing results and alignments for the hippocampal snRNA-seq study are deposited at GEO under accession code GSE314609.
- The raw MS results for the Tau interactomics study are deposited at MassIVE with the ID MSV000100283.
- This paper does not report original code.
- Any additional information required to reanalyze the data reported in this paper is available from the lead contact upon request.

## ACKNOWLEDGEMENTS

This work was supported by National Institutes of Health grants R01AG074273 (to X.C.), T32GM007752 (to L.M.R. and M.B.), T32AG066596 (to L.M.R.), F99AG088570 (to L.M.R.), and P30NS047101 (UCSD SoM Microscopy Core). D.M.M. is an awardee of an Alzheimer’s Association Research Fellowship to Promote Diversity. We thank B. Gastfriend, the San Diego Supercomputer Center Triton Shared Computing Cluster, and the Zhong Lab in UCSD’s Dept. of Bioengineering for computational assistance. We thank J. Girardini, G. Ramirez, Y. Tao, X. Lyu, Y. Liu, Y. Li, N. Athavale, and B. Bai for technical assistance and members of the Chen lab for discussions.

## AUTHOR CONTRIBUTIONS

Conceptualization, L.M.R., J.C.N., and X.C.; methodology, L.M.R., M.B., Y.N., Q.X., and X.C.; investigation, L.M.R., J.F., J.D., L.L., M.B., Y.N., D.M.M., Q.X., A.D.L., S.J., and M.R.; writing—original draft, L.M.R. and X.C.; Writing—review & editing, all authors; visualization, L.M.R., J.D., A.D.L., and X.C.; funding acquisition, L.M.R., M.B., and X.C.; resources and supervision, R.D., J.C.K., J.C.N., S.A.M., and X.C.

